# Rapidly induced drug adaptation mediates escape from BRAF inhibition in single melanoma cells

**DOI:** 10.1101/2020.03.15.992982

**Authors:** Chen Yang, Chengzhe Tian, Timothy E. Hoffman, Nicole K. Jacobsen, Sabrina L. Spencer

## Abstract

Despite increasing numbers of effective anti-cancer therapies, successful treatment is limited by the development of drug resistance. While the contribution of genetic factors to drug resistance is undeniable, little is known about how drug-sensitive cells first evade drug action to proliferate in drug. Here we track the response of thousands of single melanoma cells to BRAF inhibitors and show that a subset escapes drug within the first 3 days of treatment. Cell-cycle re-entry occurs via a non-genetic mechanism involving activation of mTORC1 and ATF4, validated in cultures of patient biopsies. These escapees cycle periodically in drug, incur significant DNA damage, and out-proliferate non-escapees over extended treatment. Our work reveals a mutagenesis-prone, expanding subpopulation of early drug escapees that may seed development of permanent drug resistance.

## Main text

Melanomas driven by the BRAF^V600E^ mutation (Fig. 1A) show immediate, positive clinical response to BRAF inhibitors vemurafenib and dabrafenib, but resistance nevertheless develops within months (*1*–*4*). A non-genetic, reversible drug-tolerant state has been reported both in the clinic and preclinical cancer models, delineating a critical step on the road to permanent resistance (*5*–*14*). However, little is known about the inception of drug tolerance, in particular the timing and mechanisms involved in cancer cell adaptation to drug.

**Fig. 1.**
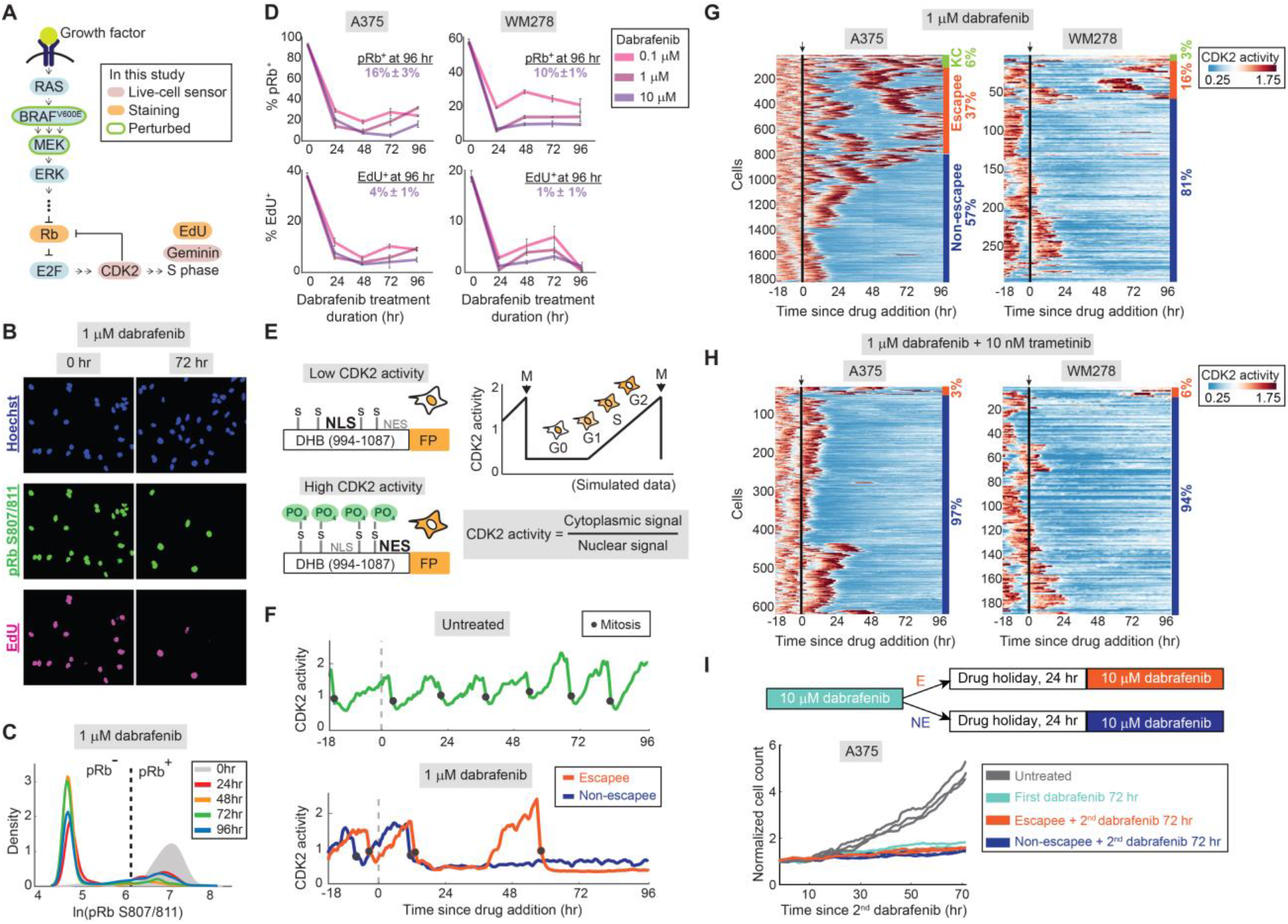
A subpopulation of melanoma cells can rapidly and reversibly escape BRAF inhibition. (**A**) Schematic diagram of MAPK-dependent cell-cycle entry. (**B**) A375 cells treated with 1 μM dabrafenib for 0 or 72 hr and stained for proliferation markers phospho-Rb and EdU. (**C**) Probability density of phospho-Rb S807/811 intensity in A375 cells. The vertical dashed line marks the saddle point between pRb^+^ and pRb^−^. (**D**) Percentage of pRb^+^ and EdU^+^ cells in A375 and WM278 cells treated for the indicated drug doses and lengths of time; % positive cells is noted for the highest dose. Error bars: mean ± std of 3 replicate wells. (**E**) Schematic of CDK2 sensor (*15*). (**F**) Representative single-cell traces of CDK2 activity in an untreated A375 cell (top), and a 1 μM dabrafenib-treated escapee and non-escapee (bottom). (**G**) Heatmap of single-cell CDK2 activity traces in 1 μM dabrafenib-treated A375 and WM278 cells. Each row represents the CDK2 activity in a single cell over time according to the colormap. Apoptotic cells (fig. S1D) are not included in the heatmap. The percentages mark the proportion of cells with each behavior. KC, keep cycling. Arrow and black line mark the time of drug addition. (**H**) Heatmap as in (G) of single-cell CDK2 activity in A375 and WM278 cells treated with dabrafenib and trametinib. (**I**) Cell count over time as measured in triplicate by time-lapse microscopy after a 24 hr drug holiday.

To examine the initial response to dabrafenib, we treated two BRAF^V600E^ melanoma lines (A375 and WM278) with dabrafenib and found that MAPK signalling and proliferation were initially repressed but rebounded within three days of treatment (fig. S1A). Single-cell immunofluorescence identified a dose-dependent subpopulation of residual proliferating cells in SKMEL19, A375, and WM278 melanoma lines (Fig. 1, B to D and fig. S1, B and C). Cell death was low during this period, consistent with the cytostatic nature of this drug (fig. S1D).

Does this residual proliferative population arise from a few initial cells and their offspring, or can many cells cycle occasionally in drug? To answer this question, we monitored cell proliferation in real time using long-term time-lapse imaging of a fluorescent biosensor for CDK2 activity (*15*) (Fig. 1E) coupled with EllipTrack (*16*), our new cell-tracking pipeline optimized for hard-to-track cancer cells. In the untreated condition, the vast majority of cells cycle rapidly and continuously (Fig. 1F and fig. S2A; Movie S1). Following drug treatment, we discovered a surprising degree of behavioral heterogeneity: while the majority of cells respond to dabrafenib by entering a quiescent CDK2^low^ state for the remainder of the imaging period (non-escapee), a subset of cells initially enters a CDK2^low^ quiescence but later escapes drug treatment by building up CDK2 activity to re-enter the cell cycle and divide (escapee) (Fig. 1F and Movie S2). Clustering thousands of single-cell drug responses in A375 and WM278 cells revealed that the majority were non-escapees, a smaller subset were escapees, and a few percent kept cycling, albeit more slowly (Fig. 1G and fig. S2B), consistent with a dose-dependent increase in cell count over time (fig. S2C). Co-treatment of dabrafenib with trametinib, a clinically approved MEK1/2 inhibitor shown to block MAPK pathway reactivation in melanoma cells treated with BRAF inhibitor (*5*), reduced the escapee phenotype but did not fully eliminate it (Fig. 1H and fig. S2, D and E; Movies S3 and S4).

In testing characteristics that enable drug escape, we noted that cell-cycle phase at the time of drug addition did not influence the potential to escape (fig. S2, F and G), nor did the existence of a small subpopulation of naturally quiescent cells (fig. S2H). To determine if pre-existing drug-resistance mutations were at the root of rapid escape, we examined whether escapees could revert to a drug-sensitive state after drug withdrawal. We used mCerulean-Geminin (*17*), a protein that accumulates in S/G2 and is absent during G0/G1, to identify escapees (Geminin^+^) and non-escapees (Geminin^−^) (fig. S2I). Cells were treated with dabrafenib for 72 hr and sorted into Geminin^+^ and Geminin^−^populations, given a 24 hr drug-free holiday, and then were filmed upon dabrafenib re-treatment. If the ability to escape from dabrafenib results from pre-existing drug-resistance mutations, the proliferation rate of sorted escapees during the second round of dabrafenib treatment should be significantly higher than that of sorted non-escapees and drug-naïve cells (fig. S2I). Instead, we observed that the proliferation rate was indistinguishable among these three populations (Fig. 1I), indicating that the ability to escape from dabrafenib during the first few days of treatment is not due to pre-existing resistance mutations but rather to a reversible cellular rewiring.

Since we failed to identify pre-existing cell states associated with escapees, and since escapees could not be eliminated by dabrafenib/trametinib combination therapy, adaptive mechanisms in addition to MAPK pathway reactivation must exist in escapees. We therefore performed scRNA-seq on A375 cells treated with dabrafenib for 72 hr (fig. S3A). Escapees were identified by computing the proliferation probability based on expression of 51 cell-cycle genes (Table S1), two of which were validated by RNA fluorescence in situ hybridization (RNA FISH; fig. S3, B and C). Cells with a proliferation probability of one were classified as “high-confidence proliferative cells” and cells with probabilities lower than e^−40^ as “high-confidence quiescent cells” (fig. S3D), which can be visualized on a t-SNE plot (Fig. 2A). Escapees can then be identified as a small peninsula within the treated population that points downward toward the untreated population (Fig. 2A).

**Fig. 2.**
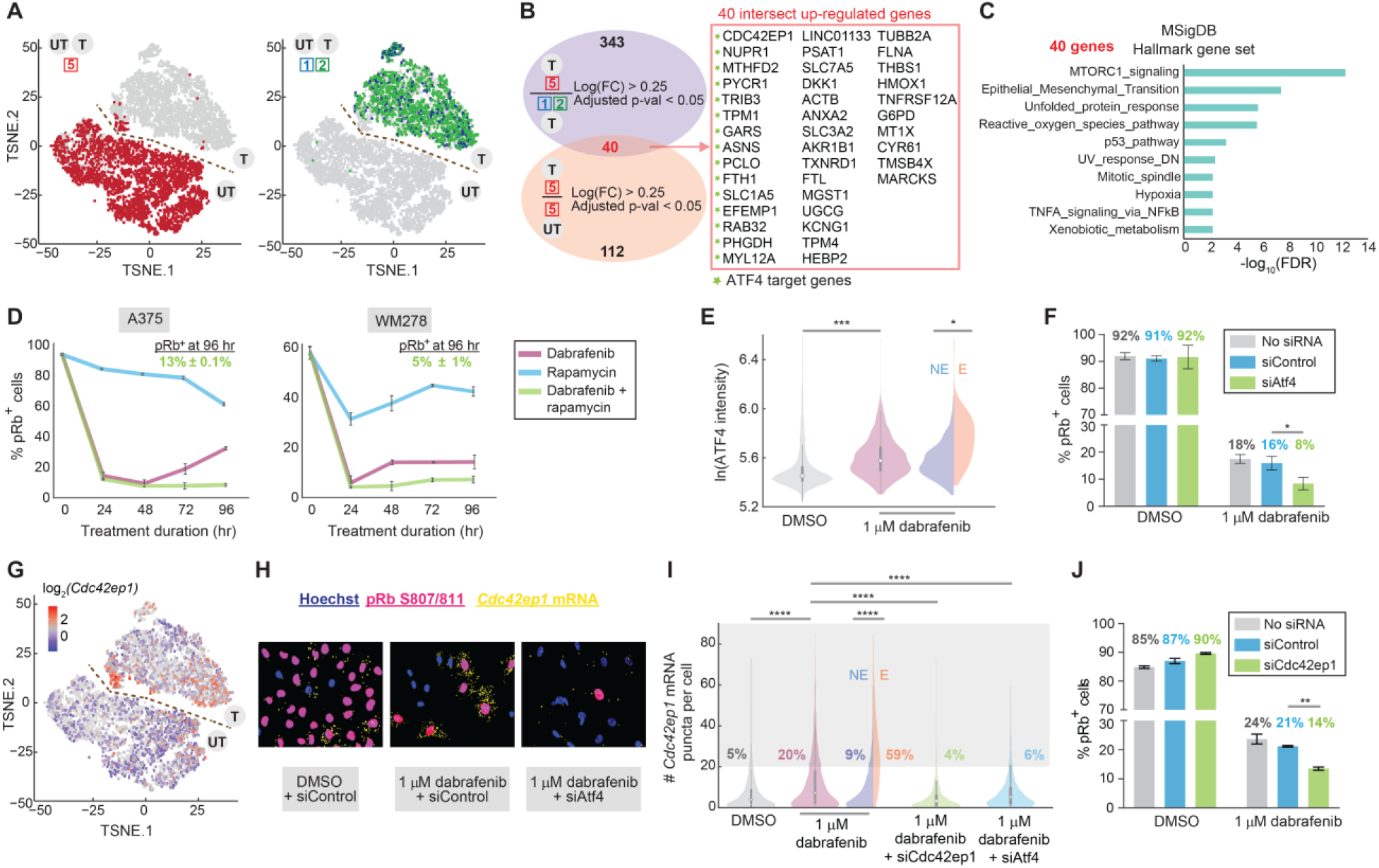
scRNA-seq reveals new gene targets associated with escape from dabrafenib. (**A**) Co-visualization of untreated and treated scRNA-seq datasets on a single t-SNE plot, showing high-confidence proliferative cells in red and high-confidence quiescent cells in blue and green (see fig. S3D for definition of boxed numbers). Escapees can be identified as a small peninsula of red proliferative cells in the treated condition. (**B**) Venn diagram of differentially expressed genes as described in the text. (**C**) MSigDB hallmark gene set enrichment analysis of the 40 genes using the false-discovery rate cutoff of 0.05. (**D**) Percentage of pRb^+^ cells in A375 or WM278 cells treated for indicated durations with 1μM dabrafenib or 10 nM rapamycin alone, or in combination. Error bars: mean ± std of 3 replicate wells. (**E**) Violin plot showing ATF4 protein levels by immunofluorescence in A375 cells. Split violin shows ATF4 levels in non-escapees (NE) and escapees (E) identified by phospho-Rb (S780) co-staining. Each population value is pooled from 3 replicate wells. (**F**) The percentage of pRb^+^ cells in the indicated conditions. Error bars: mean ± std of 4 replicate wells. (**G**) Visualization of single-cell *Cdc42ep1* mRNA expression levels on the combined t-SNE plot showing increased expression in escapees. (**H**) Representative RNA-FISH images for *Cdc42ep1* with phospho-Rb (S807/811) and Hoechst staining. (**I**) Violin plot showing the number of mRNA puncta for *Cdc42ep1* in the indicated conditions. The percentage of cells with > 20 mRNA puncta is indicated on the plot. Each population value is pooled from 2 replicate wells. (**J**) The percentage of pRb^+^ cells in the indicated conditions. Error bars: mean ± std of 4 replicate wells.

We first assessed gene expression at a population level to relate our data to the “AXL/MITF phenotype-switch” model derived from bulk RNA-seq data, where cells can either adopt a differentiated AXL^low^/MITF^high^ or a dedifferentiated AXL^high^/MITF^low^ phenotype (*18*–*20*). On average, dabrafenib-treated A375 cells adopt an AXL^low^/MITF^high^ gene-expression state (fig. S4, A to C), consistent with observations that melanoma patients treated with MAPK inhibitors initially show an increase in MITF (*8*). In contrast, at the single-cell level, the escapee subpopulation exists in an AXL^high^/MITF^low^ state (*21*–*23*) (fig. S4, D and E). Dedifferentiation is often mediated by SOX10 loss (*18*, *24*, *25*) and consistently, we saw lower expression of SOX10 in escapees (fig. S4, A and B). NGFR, another melanoma drug-resistance marker whose expression marks neural crest stem cells (*20*, *22*, *25*), was induced in the treated population, though no significant difference was observed between escapees and non-escapees (fig. S4, A to C and fig. S4E). Thus, while the treated population on average appears to be in an AXL^low^/MITF^high^ state, the escapee subpopulation is in a more dedifferentiated AXL^high^/MITF^low^ state. We therefore propose that escapees could be the seed population that drives the drug-induced melanoma dedifferentiation signature.

To identify new genes and pathways involved in escape from dabrafenib, we derived a list of genes that were differentially expressed in both dabrafenib-treated escapees *vs*. non-escapees and in dabrafenib-treated escapees *vs*. untreated proliferating cells. This yielded 40 upregulated genes and 16 downregulated genes (Fig. 2B and Table S2). *Cis*-regulatory sequence analysis in iRegulon revealed 15 transcriptional targets of ATF4 among the 40 upregulated genes. ATF4 is induced by the Integrated Stress Response, which impairs general translation but enhances translation of ATF4, leading to upregulation of a group of stress-responsive genes (*26*). Interestingly, ATF4 was reported to maintain an AXL^high^/MITF^low^ phenotype in melanoma (*27*), consistent with our findings. Pathway enrichment analysis with the Hallmark Gene Set identified mTORC1 signalling as enriched in escapees, in addition to other stress-response signatures such as unfolded protein response, oxidative stress, and p53-dependent pathways (Fig. 2C). Additionally, the epithelial-mesenchymal transition pathway was enriched, consistent with escapees possessing a mesenchymal-like dedifferentiated gene signature.

Activation of mTORC1 and ATF4 pathways in escape from BRAF inhibition illuminates potential new targets to extinguish the escapee population. Indeed, treatment with dabrafenib plus mTORC1 inhibitor rapamycin blocked the rebound of escapees normally seen with dabrafenib alone (Fig. 2D). In addition, ATF4 mRNA and protein were upregulated upon dabrafenib treatment, with protein levels particularly high in escapees (Fig. 2E and fig. S5A). Dabrafenib-mediated ATF4 upregulation could be reduced upon co-treatment with rapamycin, suggesting that mTORC1 and ATF4 activities are coupled (fig. S5B). If ATF4 activation promotes escape from dabrafenib, then its depletion should also reduce the percentage of escapees. Indeed, siRNA knockdown of ATF4 reduced escapees by 50% relative to control siRNA (Fig. 2F).

As the top hit among ATF4 target genes, *Cdc42ep1*, a member of the Rho GTPase family (*28*), represents a candidate that may mediate escape (Fig. 2G). Another induced ATF4 target gene of interest is *Rab32* (fig. S5C), belonging to a family of Ras-related GTPases (*29*). The mRNA expression of both genes was enriched in escapees upon dabrafenib treatment and markedly decreased after ATF4 knockdown, confirming their ATF4-target status (Fig. 2, H and I and fig. S5, D and E). Knockdown of *Cdc42ep1* or *Rab32* in dabrafenib-treated cells significantly reduced the percentage of escapees compared to siControl (Fig. 2J and fig. S5F). Similar results were obtained for these two genes in WM278 cells (fig. S5, G to I). *Linc01133*, a lncRNA of unknown function and the top hit among non-ATF4 target genes, was also significantly enriched in escapees (fig. S5, J and K) and enabled dabrafenib-mediated escape (fig. S5L), indicating that a fraction of escapees relies on ATF4-independent mechanisms to proliferate in drug.

To probe the translational relevance of our findings, we tested for the existence of escapees in two short-term *ex vivo* cultures of BRAF^V600E^ melanoma patient biopsies obtained prior to treatment (MB4562 and MB3883). Dose response curves revealed residual cycling cells even at the maximal dose of dabrafenib (Fig. 3A), indicating presence of escapees in these patient samples. Using the IC50 for each patient sample, we observed a decrease in cycling cells at 4 days of treatment followed by a significant increase at 7 days of treatment, similar to the rebound in proliferation observed in commercial lines (Fig. 3B). We then measured both ATF4 and phospho-S6, a marker for mTORC1 activity, in the more drug-sensitive patient sample MB3883, and found that both signals steadily increased throughout a week of treatment, with significant enrichment in escapees (Fig. 3, C and D). Consistent with the observed ATF4 induction, *Cdc42ep1* and *Rab32* mRNA levels were also induced in dabrafenib-treated MB3883 cells with significant enrichment in escapees relative to non-escapees (Fig. 3, E and F and fig. S6, A and B).

**Fig. 3.**
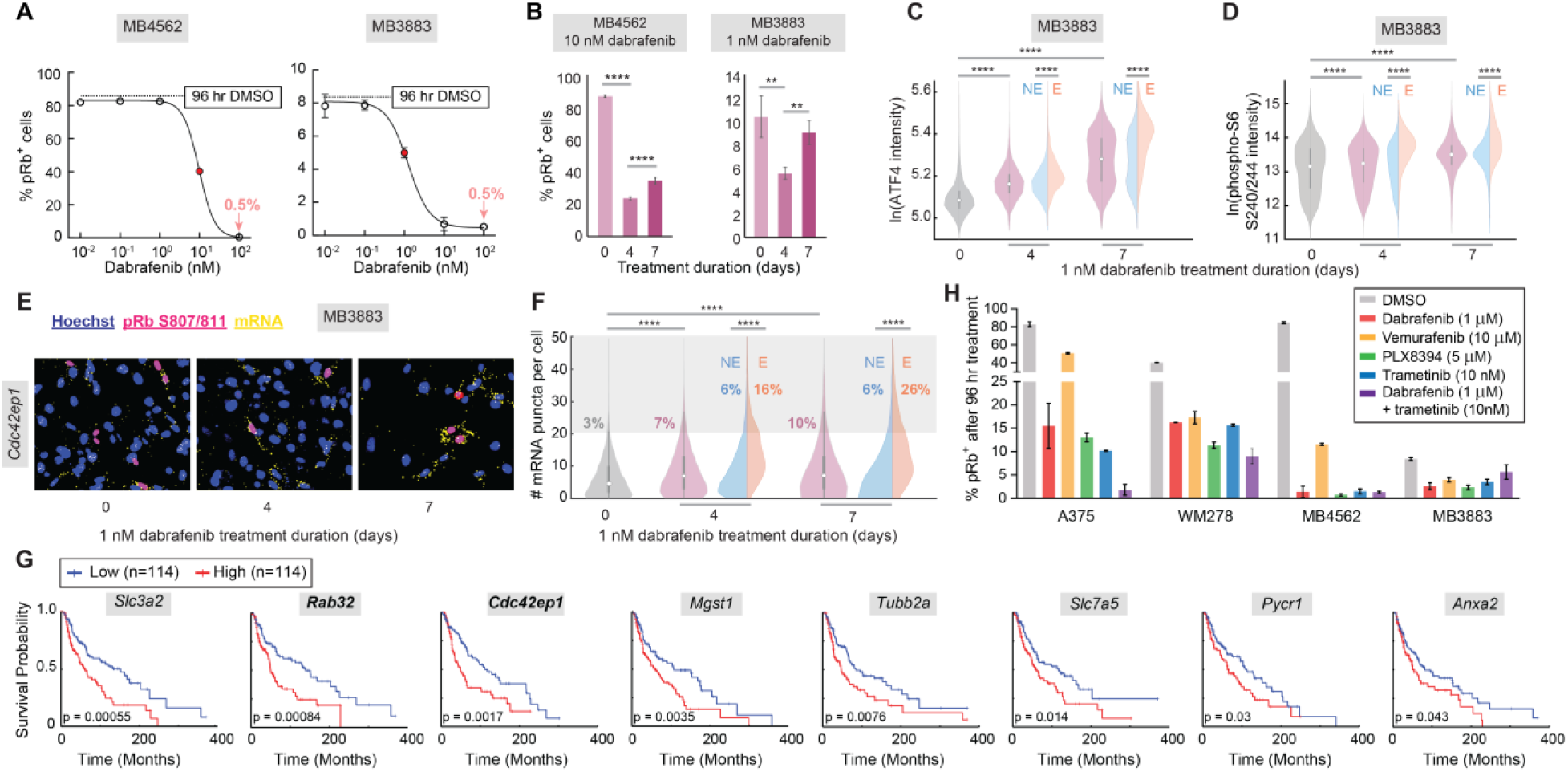
Existence of escapees and upregulation of ATF4 target genes in clinical samples. (**A**) Dose response curves at 96 hr of treatment in two *ex vivo* patient cultures, noting a residual 0.5% of pRb^+^ cells at the highest dose. IC50 values used in further experiments are displayed as red dots. Error bars: mean ± std of 3 replicate wells. (**B**) Percentage of pRb^+^ cells in the two patient samples treated with IC50 dose of dabrafenib for 4 or 7 days. Error bars: mean ± std of 3 replicate wells. (**C** and **D**) Violin plots showing ATF4 and phospho-S6 (S240/244) levels by immunofluorescence in MB3883 patient cells treated with 1 nM dabrafenib for 0, 4, or 7 days. Escapees identified by phospho-Rb (S780) or EdU co-staining for ATF4 or phospho-S6, respectively. Each population value is pooled from 6 replicate wells. (**E**) RNA-FISH images for *Cdc42ep1* with pRb (S807/811) and Hoechst staining, for MB3883 cells cultured in 1 nM dabrafenib for 0, 4, or 7 days. (**F**) Violin plot showing the number of *Cdc42ep1* mRNA puncta in MB3883 cells. The percentage of cells with > 20 mRNA puncta is indicated on the plot. Each population value is pooled from 2 replicate wells. (**G**) Melanoma patient survival curves for 8 of 40 genes upregulated in escapees. (**H**) Percentage of pRb^+^ cells in A375, WM278, and two *ex vivo* patient cultures treated with high doses of dabrafenib, vemurafenib, PLX8394, trametinib, or dabrafenib plus trametinib for 4 days. Error bars: mean ± std of at least 2 replicate wells.

To determine the long-term clinical relevance of these results, we assessed our list of 40 genes uniquely upregulated in A375 escapees for impact on melanoma patient survival using The Cancer Genomics Atlas (TCGA). Eight out of the 40 genes were associated with significantly worse survival, including *Cdc42ep1*, *Rab32,* and several other ATF4 and mTORC1-associated targets (Fig. 3G), representing a strikingly high proportion of the 40 genes (*p* = 4e-9, Chi-square test). Thus, genes uniquely upregulated in A375 escapees after just days of dabrafenib treatment show negative long-term impact on patient survival assessed over decades.

Do these findings pertain only to dabrafenib treatment, or do they extend to other MAPK pathway inhibitors? We treated two commercial lines (A375 and WM278) and the two *ex vivo* patient biopsies (MB4562 and MB3883) with several BRAF inhibitors (dabrafenib, vemurafenib, PLX 8394) and a MEK1/2 inhibitor (trametinib). Cycling cells could not be completely eliminated by any of the treatments, even at high drug doses (Fig. 3H). Additionally, ATF4, *Cdc42ep1*, and *Rab32* levels were specifically enriched in escapees in every treatment condition (fig. S6, C to E). These findings suggest that the adaptive escape phenomenon and induction of the ATF4 stress response may be widespread among different modes of MAPK pathway inhibition and may also be clinically relevant.

Could escapees be the seed population driving eventual acquisition of drug-resistance mutations? For this to be the case, escapees would have to be both prone to mutagenesis and out-proliferate non-escapees. Indeed, A375 and MB3883 escapees showed increased γ-H2AX straining and increased double-strand breaks relative to non-escapees (Fig. 4, A and B and fig. S7, A and B). Furthermore, dabrafenib-treated EdU^+^ cells reach a lower EdU intensity maximum compared with untreated cycling cells (fig. S7, C and D), suggesting a reduced DNA synthesis rate. Suspecting stalled DNA replication forks, we stained cells for FANCD2, a protein that localizes to stalled replication forks (*30*). FANCD2 staining was elevated in drug-treated cells and was particularly high in EdU^+^ escapees undergoing DNA replication (Fig. 4C). Another potential cause of reduced DNA synthesis rate is the under-licensing of origins of replication prior to the start of S phase. We therefore measured the amount of the licensing factor MCM2 bound to chromatin in single cells (*31*) and found marked under-licensing after dabrafenib and trametinib treatment (Fig. 4D and fig. S7, E to G). Thus, escapees experience dysregulated licensing and heightened DNA replication stress, which are known to cause genomic instability in cancer cells (*32*). Together, these data demonstrate that cells cycling in the presence of dabrafenib are prone to mutagenesis.

**Fig. 4.**
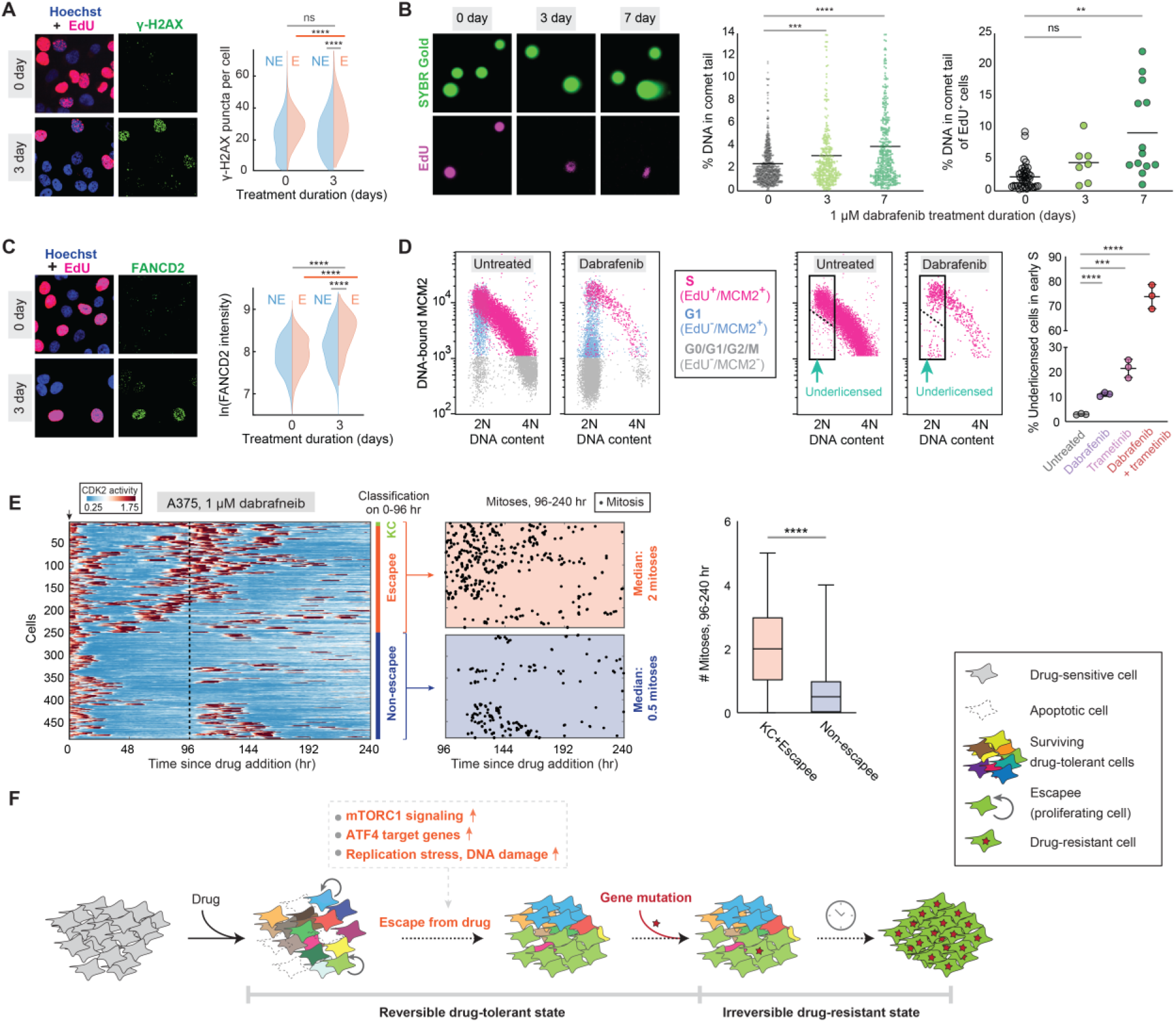
Escapees are prone to DNA damage and outgrow non-escapees over extended drug treatment. (**A**) Images of 1 μM dabrafenib-treated A375 cells stained for EdU and γ-H2AX. Quantification of γ-H2AX puncta for escapees and non-escapees is plotted as split violins. Each population value is pooled from 4 replicate wells. (**B**) Neutral comet assay in dabrafenib-treated A375 cells. Gels were co-stained for EdU incorporation. Plots indicate percent tail intensity over each entire comet, with mean values displayed as a horizontal line. Each population value is pooled from 2 biological comet slide replicates. (**C**) Images of 1 μM dabrafenib-treated A375 cells stained for EdU and FANCD2. Quantification of FANCD2 intensity for escapees and non-escapees is plotted as split violins. Each population value is pooled from 4 replicate wells. (**D**) Flow cytometric analysis of DNA replication licensing determined by chromatin-bound MCM2 in A375 cells treated with BRAF and/or MEK inhibitors. Cells were first gated on EdU incorporation (fig. S7F) and DNA-bound MCM2 was then plotted relative to DNA content. Right-most plot shows percent under-licensed cells relative to all early S phase cells (cells under dashed line / all cells in rectangle); mean ± std of 3 replicate samples. (**E**) CDK2 activity heatmap for 500 dabrafenib-treated single cells tracked over 10 days. Cell behavior was classified based on the first 96 hr, and subsequent mitosis events between 96 and 240 hr are plotted as black dots or as a boxplot. (**F**) Model schematic as described in the text.

To determine whether escapees out-proliferate non-escapees in the longer term, we imaged and tracked dabrafenib-treated A375 cells over 10 days. Cells were classified as escapees or non-escapees based on behavior in the first 96 hr, and their proliferative activity was examined during the remaining 6 days. Non-escapees rarely re-entered the cell cycle during the final 6 days, with a median of 0.5 mitoses. By contrast, escapees cycled significantly more frequently than non-escapees, having a median of two mitoses with long quiescence periods in between (Fig. 4E and fig. S8). Thus, despite incurring DNA damage, escapees out-proliferate non-escapees and will dominate the population in the long term.

In summary, we discovered that a subset of melanoma cells can rapidly adapt to drug treatment and proliferate via activation of alternative signalling pathways. Because escapees are prone to DNA damage and yet out-proliferate non-escapees, they may represent a seed population enabling permanent (genetic) drug resistance (Fig. 4F). Since none of the clinically approved MAPK pathway inhibitors tested here alone or in combination successfully block proliferation in 100% of commercial cells or primary patient samples, these findings could have broad applicability, implying that non-genetic escape from targeted therapies may be much more common than currently appreciated. New drug combinations targeting mTORC1, ATF4, or the adaptation state more broadly, could forestall drug resistance and tumor relapse.

## Acknowledgements

We thank Xuedong Liu, Kasey Couts, Grace Zheng, and Rebecca Schweppe for comments on the manuscript; members of Spencer lab for general help and discussion; Theresa Nahreini and the cell culture facility for cell sorting; Joseph Dragavon at the BioFrontiers advanced light microscopy core; Kasey Couts and William Robinson at the University of Colorado Skin Cancer Biorepository. This work was conducted with the help and resources of the BioFrontiers Computing Core at the University of Colorado BioFrontiers Institute.

## Funding

The PerkinElmer Opera Phenix is supported by NIH grant 1S10ODO25072. The Aria Fusion FACS sorter and BD FACSCelesta are supported by NIH grant S10ODO21601. This work was supported by a Kimmel Scholar Award (SKF16-126), a Searle Scholar Award (SSP-2016-1533), and a Beckman Young Investigator Award to S.L.S.

## Author Contributions

C.Y. conducted the majority of experiments, analyses, data interpretation, and manuscript preparation; C.T. developed EllipTrack, helped analyse microscopy and scRNA-seq data and assisted with manuscript preparation; T.E.H. assisted with experiments, analyses, and manuscript preparation. C.Y., C.T., T.E.H. and N.K.J. manually verified cell traces; S.L.S. conceived the project, suggested the experiments, interpreted the data, and wrote the manuscript with C.Y.

## Competing interests

The authors declare no competing interests.

## Data and materials availability

Further information and requests for resources and reagents should be directed to and will be fulfilled by the Lead Contact, Sabrina Spencer.

## Materials and Methods

### Cell culture and cell line generation

The A375 melanoma cell line (#CRL-1619) was purchased from American Type Culture Collection (ATCC). A375 cells were cultured at 37 °C with 5% CO_2_ in DMEM (Thermo Fisher, #12800-082) supplemented with 10% FBS, 1.5g/L sodium bicarbonate (Fisher Chemical, #S233-500), and 1X penicillin/streptomycin. The WM278 and SKMEL19 cell lines were obtained from Dr. Natalie Ahn (University of Colorado Boulder) and Dr. Neal Rosen (Memorial Sloan Kettering Cancer Center), respectively. WM278 and SKMEL19 cells were maintained at 37 °C with 5% CO_2_ in RPMI1640 (Thermo Fisher, #22400-089) supplemented with 10% FBS, 1X Glutamax, 1X sodium pyruvate (Thermo Fisher, #11360-070), and 1X penicillin/streptomycin. The A375 and WM278 lines were submitted for short tandem repeat (STR) profiling and were authenticated as exact matches to A375 and WM278, respectively. MB3883 and MB4562 cells were obtained from the Cutaneous Oncology Melanoma Bank at the University of Colorado and were maintained in the same conditions as WM278 and SKMEL19. At the time the patient cells were received, the MB3883 line had been passed through PDX models, whereas the MB4562 line was directly cultured following initial patient biopsy. A375 and WM278 cells were transduced with H2B-mIFP and DHB-mCherry lentivirus or with H2B-mCherry and mCerulean-Geminin lentivirus as described previously (*15*). Cells stably expressing these sensors were isolated by two rounds of FACS.

### Small molecules

Drugs used in this study are: dabrafenib (Selleckchem, #S2807), trametinib (Selleckchem, #2673), vemurafenib (Selleckchem, #S1267), PLX8394 (MedChemExpress, HY-18972), rapamycin (Selleckchem, #S1039), and etoposide (Selleckchem, #S1225).

### Antibodies

The antibodies used for this study are: phospho-ERK T202/Y204 (1:1000) (Cell Signaling Technology, #4370), phospho-Rb S807/811 (1:1000) (Cell Signaling Technology, #8516P), GAPDH (1:2000) (Cell Signaling Technology, #5174), ATF4 (1:200) (Cell Signaling Technology, #11815S), phospho-Rb S780 (1:1500) (BD Biosciences, #558385), phospho-S6 S240/244 (1:250) (Cell Signaling Technology, #2215), AXL (1:200) (Cell Signaling Technology, #8661), MITF (1:100) (Abcam, #ab3201), NGFR (1:1000) (Cell Signaling Technology, #8238), SOX10 (1:1000) (Cell Signaling Technology, #89356), FANCD2 (1:500) (Novus Biologicals, #NB100-182), γ-H2AX (1:400) (Cell Signaling Technology, #9718), and MCM2 (1:200) BM28), (BD Biosciences, #610700). Secondary antibodies were used at 1:500 dilution (anti-rabbit Alexa Fluor-647: Thermo Fisher, #A-21245; anti-rabbit Alexa Fluor-488: #A-11034; anti-mouse Alexa Fluor-488: #11029; anti-mouse Alexa Fluor-546: #11030); and anti-rabbit IgG, HRP-linked secondary antibody (Cell Signaling Technology, #7074S) for western blotting at 1:1000.

### Western blot

Cells were washed 3 times with PBS and lysed in 2x LDS sample buffer (Thermo Fisher, #B0008) supplemented with reducing reagent and 1x phosphatase and protease inhibitor. Lysates were sheared with a 1cc U-100 insulin syringe (Becton Dickinson, #329424) and heated at 95 °C for 10 min. Proteins were separated by Bolt 4-12% Bis-Tris Plus gel (Thermo Fisher, NW04125BOX) and transferred to a PVDF membrane (Merck Millipore, #IPFL00010). The membrane was incubated in 3% BSA (GoldBio, #A-421-250) supplemented with 0.1% Tween-20 (Thermo Fisher, #9005-64-5) at room temperature for 2 hr before overnight incubation with antibodies against phospho-ERK^T202/Y204^ (1:1000), phospho-Rb^S807/811^ (1:1000), and GAPDH (1:2000). The membrane was then washed for 5 min with PBS supplemented with 0.1% Tween-20 five times and then incubated with anti-rabbit IgG, HRP-linked secondary antibody. The chemiluminescent signals were detected on an Azure C600 from Azure Biosystems.

### EdU incorporation assay

To identify cells in S phase of the cell cycle, cells were pulsed with 10 μM EdU at 37 °C for 15 minutes prior to fixation with 4% paraformaldehyde. The EdU was visualized as described in the manufacturer’s protocol (Thermo Fisher, #C10340 and #C10641). Cells were then twice washed with PBS and blocked with 3% BSA for 1 hr at room temperature to prepare for further immunostaining.

### Apoptosis assay

Cells were seeded in 12-well plates (Corning, #3513), at 10^5^ cells/well, 24 hr prior to drug treatments. Wells were treated in triplicate with various doses and combinations of MAPK pathway inhibitors used in the study for 2 days, 4 days, or 2 weeks. Etoposide (10 μM) was used as a positive control for apoptosis. After treatment, non-adherent cells were first harvested by pipetting, adherent cells were harvested by trypsinization, and these two populations were then combined. Cell suspensions were centrifuged and resuspended in calcium-rich binding buffer provided by the apoptosis staining kit (abcam, #ab14085) to reach ~10^6^ cells/mL. Live suspensions were stained with both Annexin V-FITC (1:100) and propidium iodide, PI (1:100) for 5 minutes, and were subsequently spun and washed with binding buffer to remove excess dye. Single-cell fluorescent signals were acquired on a BD FACSCelesta flow cytometer equipped with 488 and 561 nm lasers. By convention, cells were gated in FlowJo to remove debris and doublets and FlowJo’s compensation matrix was used to correct for any bleed-through between FITC and PI. Annexin V-FITC and PI values were plotted as a bivariate scatter and etoposide-determined quadrant gating was applied to all plots to reach final apoptotic population percentages.

### siRNA transfection

siRNA transfections were performed using the DharmaFECT 4 reagent (Dharmacon, #T-2004-02) according to the manufacturer’s instructions. The transfection mix was added to the cells at the time of drug treatment and removed after 6 hr. The knockdown efficiency was determined by RNA FISH 72 hr post-transfection. Oligonucleotides used in this study are: DS NC-1 (IDT, #51-01-14-04), LINC01133 DsiRNA (IDT, #hs.Ri.LINC01133.13.1, #hs.Ri.LINC01133.13.2, #hs.Ri.LINC01133.13.3), ATF4 DsiRNA (IDT, #hs.Ri.ATF4.13.3, #hs.Ri.ATF4.13.1), RAB32 DsiRNA (IDT, #hs.Ri.RAB32.13.1, #hs.Ri.RAB32.13.2, #hs.Ri.RAB32.13.3), CDC42EP1 DsiRNA (IDT, #hs.Ri.CDC42EP1.13.1, #hs.Ri.CDC42EP1.13.2, #hs.Ri.CDC42EP1.13.3)

### RNA FISH and immunofluorescence

Cells were seeded on a glass-bottom 96-well plate coated with collagen 24 hr prior to drug treatment. Cells were fixed with 4% paraformaldehyde, and when applicable, were processed for RNA FISH analysis according to the manufacturer’s protocol (ViewRNA ISH Cell Assay Kit) (Thermo Fisher, #QVC0001). mRNA probes were hybridized at 40 °C for 3 hr, followed by standard amplification and fluorescent labelling steps also at 40 °C. Probes used in this study are ViewRNA Type 6 probes from Thermo Fisher: CDC42EP1 (VA6-3170107-VC), RAB32 (VA6-3175871-VC), CCNA2 (VA6-15304-VC), CCNB1 (VA6-16942-VC), LINC01133 (VA6-20432-VC). For quantification of individual mRNA puncta, cells were stained with total protein dye, CF 568 succinimidyl ester (1:100,000) (Millipore sigma, #SCJ4600027), to create a whole-cell mask for segmentation. FISH images were taken on PerkinElmer Opera Phenix high-content screening system with a 20X 1.0 NA water objective.

For immunofluorescence, standard protocols were used: following blocking, primary antibodies were incubated overnight at 4 °C, and secondary antibodies were incubated for 2 hr at room temperature. Immunofluorescence stains without RNA FISH were imaged on a Nikon Ti-E using a 10X 0.45 numerical aperture (NA) objective.

### Live-cell imaging

Cells were seeded on a glass bottom 96-well plate coated with collagen 24 hr prior to the start of imaging. Movie images were taken on a Nikon Ti-E using a 10X 0.45 NA objective with appropriate filter sets at a frequency of 15 min per frame. Cells were maintained in a humidified incubation chamber at 37 °C with 5% CO_2_. Cells were imaged in phenol-red free full growth media (Corning, #90-013-PB) for 18 hr before treatment; the movie was then paused for drug addition and imaging continued for another 48 hr at which point the drug was refreshed; imaging then continued for another 48 hr. The drug refreshment was performed by exchanging half of the total media in each well to avoid cell loss during pipetting.

### Definition of escapees

In time-lapse microscopy experiments, escapees are defined as cells that spend more than 15 hr (one normal A375 cell cycle) or 20 hr (one normal WM278 cell cycle) in a drug-induced CDK2^low^ quiescence before building up CDK2 activity and re-entering the cell cycle. Non-escapees are defined as cells that stay in a drug-induced CDK2^low^ state until the end of the imaging period. A cell was designated a keep-cycling cell if it never entered a long quiescence (15 hr for A375 or 20 hr for WM278) during drug treatment.

In fixed-cell experiments, escapees are defined as cells that are in the cell cycle after 2 or more days of drug treatment. A cell in the cell cycle can be detected by immunofluorescence staining for phospho-Rb S807/811 or phospho-Rb S780, by EdU incorporation, by detection of mCerulean-Geminin signal, or by expression of cell-cycle genes in scRNA-seq (see Calculation of proliferation probability). We note that in all of these “snapshot” methods, escapees cannot be distinguished from keep-cycling cells since both escapees and keep-cycling cells will be engaged in the cell cycle at the time of measurement.

### FACS on escapees and non-escapees

Cells expressing mCerulean-Geminin were treated with 10 μM Dabrafenib for 72 hr before sorting on an Aria Fusion FACS machine. Untreated cells were used to choose the Geminin signal cut-off for Geminin^+^ (escapees) and Geminin^−^ (non-escapees). Drug-treated cells were directly sorted into full-growth media for 24 hr before the second round of drug treatment and the start of imaging. The subsequent imaging conditions followed the live-cell imaging protocol.

### Single-cell RNA sequencing

A375 cells were cultured with or without 1μM dabrafenib in full-growth medium for 72 hr before preparation of a single-cell suspension according to the 10X Genomics sample preparation protocol, “Single-cell suspensions from cultured cell lines for single-cell RNA sequencing”. The single GEM capture, lysis, library construction, and sequencing were performed by the Microarray and Genomics Core at the University of Colorado Anschutz Medical Campus. The untreated and the treated samples were prepared using the same chemical reagents on the same day and the libraries were sequenced in one lane of a NovaSEQ6000 with a sequencing depth of 400,000 reads/cell.

### Chromatin-bound MCM2 flow cytometry assay

Fixed-cell immunostaining of loaded MCM2 and subsequent flow cytometric analyses were performed as previously described (*31*). Briefly, following the indicated treatments, cell suspensions were fixed in 4% paraformaldehyde and washed before stepwise staining: EdU click reaction with Alexa Fluor 488 (room temperature for 30 min), mouse anti-MCM2 immunostaining (BD Biosciences #610700 at 1:200 for 1 hr at 37 °C), goat anti-mouse Alexa Fluor 546 immunostaining (Thermo Fisher #A-11003 at 1:500 for 1 hr at 37 °C), and Hoechst 33342 staining (1:10,000; overnight at 4 °C). Final cell suspensions along with staining controls were analyzed using a BD FACSCelesta flow cytometer equipped with a 405, 488, and 561 nm lasers. FCS files for each sample were captured using FACSDiva and transferred to FlowJo for analysis.

### Comet assay with EdU incorporation

The neutral comet assay was performed as described according to manufacturer protocol (Trevigen #4250-050-K) with an adaptation to capture EdU staining. Briefly, after treating cells for 30 min with EdU, cell suspensions were harvested and suspended in LM agarose gels on comet slides and processed for single-cell electrophoresis and DNA precipitation. To stain, gel sites were immersed in EdU click reaction cocktail with Alexa Fluor 647 for 30 min at room temperature. Slides were washed and then stained with SYBR Gold (Thermo Fisher, #S11494) for 30 min at room temperature. Slides were washed and dried thoroughly for 45 min before applying glycerol mountant and coverslips. Comet images were taken on a Nikon Ti-E using a 10X 0.45 numerical aperture (NA) objective. Corresponding TIFF files were processed using the ImageJ plugin OpenComet (http://www.cometbio.org/), and EdU^+^ cells were manually scored for each cell ID generated. Any doublet events and incorrect comet head segmentations were omitted from the analysis.

### Dose-response curve fitting

Dose-response curve fits for each cell line’s pRb^+^ count after dabrafenib were calculated using GraphPad Prism (v8.3). All cell line dose-response curves were fit using the standard inhibitory Hill function below. The biphasic shape of the A375 dose-response curve required the summation of two different sigmoidal inhibitory curves, both relying on the Hill equation below as well (employed in Prism as a biphasic fit function).

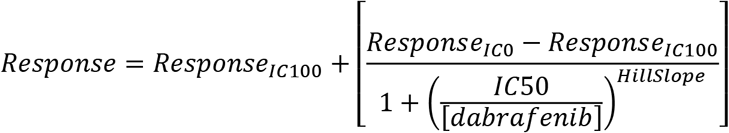

### Quantification of FISH puncta

RNA-FISH image analysis for *Cdc42ep1*, *Rab32*, and *Linc01133* was performed using Harmony high-content imaging and analysis software. First, the DAPI channel containing Hoechst DNA stain was used to identify cell nuclei in each image. Then the total protein dye CF 568 succinimidyl ester in the Cy3 channel was used to create a whole-cell mask for each cell. Cells on the border of the image were eliminated from analysis. The spot-detection function was then applied to the RNA-FISH signal in the Cy5 channel and each RNA puncta was detected as an individual spot. The mean nuclear pRb S807/811 intensity was calculated from the FITC channel. The number of RNA puncta per cell and mean intensity of phospho-Rb were then exported from Harmony software into Matlab and plotted as a violin plot of number of puncta per cell. For the high-abundance *Ccna2* and *Ccnb1* mRNAs, we used a different approach in which we measured the mean mRNA intensity in a 4-pixel ring around the nucleus.

### Statistical tests

Statistical tests were performed using GraphPad Prism. For statistical differences in single-cell immunofluorescence and FISH marker measurements represented as violin plots, *p*-values were calculated using an unpaired *t*-test with Welch’s correction for unequal variances between sample groups of all individual cells. For the number of mitoses quantified in the 10-day movie, the *p*-value was calculated using a non-parametric Mann-Whitney rank test. For bar plots representing phospho-Rb^+^ cell percentages among replicates and for differences between origin licensing measurement replicates, *p*-values were calculated using a standard unpaired *t*-test. Significance levels are reported as *p*-values less than 0.05 (*), 0.01 (**), 0.001 (***), and 0.0001 (****). Throughout the manuscript, ‘ns’ denotes no statistical significance.

### Image processing and cell tracking for time-lapse movies

Cells were tracked using EllipTrack (*16*). In brief, EllipTrack segments cells by fitting nuclear contours with ellipses. EllipTrack then utilizes a machine learning algorithm to predict cell behaviors and maps ellipses between frames by maximizing the probability of cell lineage trees. Next, signals from each color channel are extracted in the cell nuclei and cytoplasmic rings. Cell tracks were manually verified such that only cells correctly tracked during the entire movie were kept for downstream analysis. CDK2 activity was read out as the cytoplasmic:nuclear ratio of the DHB signal, as previously described (*15*). Cell count over time was determined by the number of nuclei in the field of view at each time point.

### Analysis of single-cell CDK2 traces

A customized script was used to determine whether a cell was proliferative or quiescent at each time point. In brief, we first identified “seed regions” for proliferation, which were defined as the sets of continuous time points with CDK2 activity greater than 0.9. Then, for each “seed region”, we searched the start (Restriction Point) and the end (mitosis point) of proliferation by examining the time points before and after the seed region, respectively. The Restriction Point was defined as the closest time point before the seed region with a slope of CDK2 activity less than 0.01. The mitosis point was defined as the closest time point after the seed region with a locally maximal H2B intensity. All time points between the Restriction Point and the mitosis point were assigned as proliferative, and the remaining time points were assigned as quiescent. The threshold for seed regions dropped to 0.6 for the final frames of the traces in order to identify cells that re-entered cell cycle but had yet to reach high CDK2 activity before movie ended. Finally, because the 0.9 threshold is quite high (corresponding to the time of S-phase entry), the algorithm might identify a part of G1 phase as quiescence if a cell is rapidly cycling, as in movies of untreated cells. We therefore converted all quiescence periods shorter than 4 hr to proliferative to minimize misclassification, while maintaining high accuracy in classifying drug-treated cells.

For heatmaps, escapees, non-escapees, and keep-cycling cells were plotted separately and cells within each category were sorted by the similarity of their CDK2 traces with hierarchical clustering. The plots were then combined. Apoptotic cells are not included in the heatmaps. For heatmaps sorted on cell-cycle phase, cells from all categories were aggregated and sorted by the time of their first mitosis.

### Calculation of proliferation probability

We computed the proliferation probabilities based on whether the mRNAs of 51 cell-cycle genes (Extended Data Table 1) were detected or not. Cell-cycle genes were selected from CycleBase 3.0 database (*33*) to include signature genes of all cell-cycle phases. To compute the probability of proliferation, denote the fraction of proliferative cells in the population as *p*_0_, determined experimentally by immunofluorescence staining of pRb S807/711. The values for untreated and dabrafenib-treated conditions were 0.95 and 0.18, respectively. Denote the probability of detecting at least one mRNA copy of cell-cycle gene *i* in a proliferative cell as *p*_*i*_. *p*_*i*_ was computed by dividing the fraction of cells with at least one mRNA copy detected by *p*_0_, and the value was kept between 0.01 and 0.99. Furthermore, assume the probability of detection in a quiescent cell is *ϵ*=0.01 (same for all genes). Assuming that mRNAs of different genes were independently detected, we have

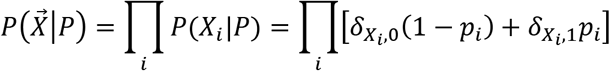

and

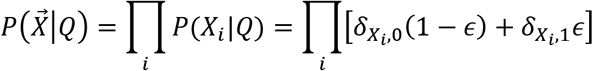

Here, P and Q stand for “Proliferative” and “Quiescent”; *X*_*i*_ is an indicator variable for detecting at least one mRNA copy of cell-cycle marker gene *i*; 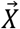 is the vector of indicator variables 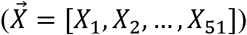; and *δ* is the Kronecker delta function. Using the fraction of proliferative cells in the population (*p*_0_) as the prior probability, and using Bayes’ theorem, we computed the probability of a cell being proliferative by

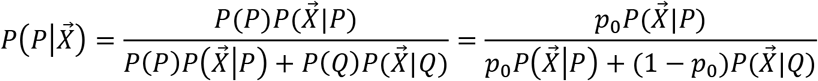

### Single-cell RNA sequencing analysis

Gene expression matrices from the untreated and treated conditions were computed by Cell Ranger (10X Genomics) and combined into a single dataset by ‘cellranger aggr’. The deeper-sequenced condition was randomly down-sampled such that two conditions had the same population-level library size in the combined dataset. Quality control was then performed such that invalid genes and outlier cells were removed(*34*). Here, a gene was denoted as invalid if it was not detected in at least one cell in both conditions, or if its gene symbol corresponded to multiple Ensemble IDs. A cell was denoted as an outlier if its log-library size or its log-number of detected genes were at least 5 median absolute deviation (MAD) lower than the population median, or if its percentage of expression mapped to mitochondrial genes was at least 5 MAD greater than the population median.

Proliferation probabilities of cells were computed as described above. Cells were classified into five groups based on their probabilities: subgroup 1, 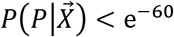; subgroup 2, 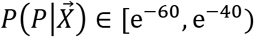; subgroup 3, 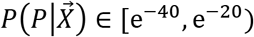, subgroup 4: 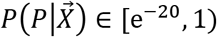; and subgroup 5, 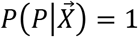. Cells belonging to subgroup 5 were denoted high-confidence proliferative cells (HC-PCs) and cells belonging to subgroup 1 and 2 were denoted high-confidence quiescent cells (HC-QCs).

Differential gene expression analysis was performed in Seurat (*35*, *36*). In brief, the gene expression matrix was normalized (method: ‘LogNormalize’, scaling factor: 10000) and scaled. The top 2000 highly variable genes were then detected (method: ‘vst’) and used for dimension reduction (Principle Component Analysis). The top 15 Principle Components were used for clustering (Louvain algorithm) (*37*) and computation of coordinates on the t-SNE plot. Finally, differentially expressed genes (DEGs) for HC-PC_T vs HC-QC_T and for HC-PC_T vs HC-PC_UT were computed (Wilcoxon Rank Sum test, logfc.threshold: 0.25, min.pct: 0.01, adjusted p-value < 0.05).

The list of 40 intersecting genes was computed by intersecting the list of upregulated DEGs for HC-PC_T vs HC-QC_T and the list of upregulated DEGs for HC-PC_T vs HC-PC_UT. ATF4 targets were detected with the iRegulon plugin (*38*) in Cytoscape (*39*). The overlap of the 40 intersecting genes with the Hallmark gene set (*40*) was computed on the Molecular Signatures Database website (*41*) (MSigDB).

Genes upregulated in escapees were defined as the upregulated (avg_logFC > 0) DEGs for HC-PC_T vs HC-QC_T, and the genes upregulated in non-escapees were defined as the downregulated (avg_logFC < 0) DEGs for HC-PC_T vs HC-QC_T. The overlap with previously published gene signatures was computed with the R package ‘GeneOverlap’ (*42*).

### Kaplan-Meier plots

Kaplan-Meier plots were generated by OncoLnc (*43*). In brief, the clinical data (as of January 5^th^, 2016) and mRNA data (Normalized RSEM values of Tier 3 RNASeqV2) were retrieved from the Skin Cutaneous Melanoma (SKCM) Project of The Cancer Genome Atlas (TCGA) (The Cancer Genome Atlas Network, 2015). Patients (n=459) with either a sequenced primary solid tumor sample (type ‘01’) or a metastatic tumor sample (type ‘06’) were used for the analysis. For patients with multiple samples sequenced, the expression levels of all samples were averaged. For each gene, patients with top and bottom 25% expression levels were used to compute Kaplan-Meier plots. Only genes with a logrank p-value less than 0.05 are included in Fig. 3.

To evaluate the enrichment of genes that had negative long-term impact on patient survival in our 40-gene list, we generated Kaplan-Meier plots for all genes in the genome as described in the paragraph above and calculated the fractions of genes that had positive and negative impacts on patient survival. Here, a gene was designated as having a positive impact if its logrank p-value was less than 0.05 and the median survival time was higher in patients with top 25% expression levels than in patients with bottom 25% expression levels; and was designated as having a negative impact if its logrank p-value was less than 0.05 and the median survival time was lower in patients with top 25% expression levels. Overall, we found that 15% and 11% of genes in the genome had positive and negative impacts, respectively. Meanwhile, 8 of the 40 genes upregulated in escapees (20%, Fig. 3) were found to have a negative impact and only one gene (NUPR1, 2.5%, data not shown) was found to have a positive impact. This contrast shows that genes with long-term negative impact on patient survival were indeed significantly enriched in our list of 40 genes upregulated in escapees (Chi-square test, p=4e-9).

### Data and code availability

All raw datasets and analysis scripts are available upon request.

---

**Table S1.** 51 cell-cycle genes used to calculate the probability of being proliferative for each cell.

|  |  |  |  |  |  |
| --- | --- | --- | --- | --- | --- |
| ARHGEF39 | CCNE1 | CENPA | KIF20B | NUF2 | UBE2C |
| ATAD2 | CCNE2 | CENPE | KIF23 | NUSAP1 |  |
| AURKB | CCNF | E2F1 | KIF4A | PLK1 |  |
| BIRC5 | CDC20 | ESCO2 | MAD2L1 | PRC1 |  |
| BRCA1 | CDC25A | FAM83D | MCM2 | RFC3 |  |
| BUB1 | CDC45 | GMNN | MCM4 | RFC4 |  |
| BUB1B | CDC6 | GTSE1 | MCM5 | RRM2 |  |
| CCNA2 | CDK1 | HJURP | MCM6 | TACC3 |  |
| CCNB1 | CDK2 | HMMR | MKI67 | TOP2A |  |
| CCNB2 | CDKN3 | KIF11 | MYBL2 | TPX2 |  |

**Table S2.**

List of 40 significantly upregulated genes and 16 significantly downregulated genes in escapees.

## Movie S1

**CDK2 activity in untreated A375 melanoma cells.** A375 cells expressing DHB-mCherry were cultured in phenol-red free full growth media. The CDK2 activity trace for the cell with the yellow arrow is plotted underneath. Red dots mark mitosis.

## Movie S2

**CDK2 activity in 10 μM dabrafenib-treated A375 cells.** A375 cells expressing DHB-mCherry were treated with 10 μM dabrafenib at the start of the movie. Two different cell behaviors (escapee and non-escapee) are sequentially displayed in this movie: the cell with a yellow arrow is a non-escapee and the corresponding CDK2 activity trace is plotted below in black; the cell with the red arrow is an escapee and the corresponding CDK2 activity trace is plotted below in red. Red dots mark mitosis.

## Movie S3

**CDK2 activity in monotherapy or combination therapy-treated A375 cells.** A) Untreated A375 cells; B) 1 μM dabrafenib-treated A375 cells. C) 10 nM trametinib-treated A375 cells. D) A375 cells treated with 1 μM dabrafenib plus 10nM trametinib. Cells were imaged in full-growth media for 18 hr before drug addition; the movie was then paused for drug addition and imaging continued for 96 hr with a pause at 48hr for drug refreshment. Flash indicates time of drug addition.

## Movie S4

**CDK2 activity in monotherapy or combination therapy-treated WM278 cells.** A) Untreated MW278 cells; B) 1 μM dabrafenib-treated WM278 cells. C) 10 nM trametinib-treated WM278 cells. D) WM278 cells treated with 1 μM dabrafenib plus 10 nM trametinib. Cells were imaged in full-growth media for 18 hr before drug addition; the movie was then paused for drug addition and imaging continued for 96 hr with a pause at 48hr for drug refreshment. Flash indicates time of drug addition.

---

**Fig. S1.**
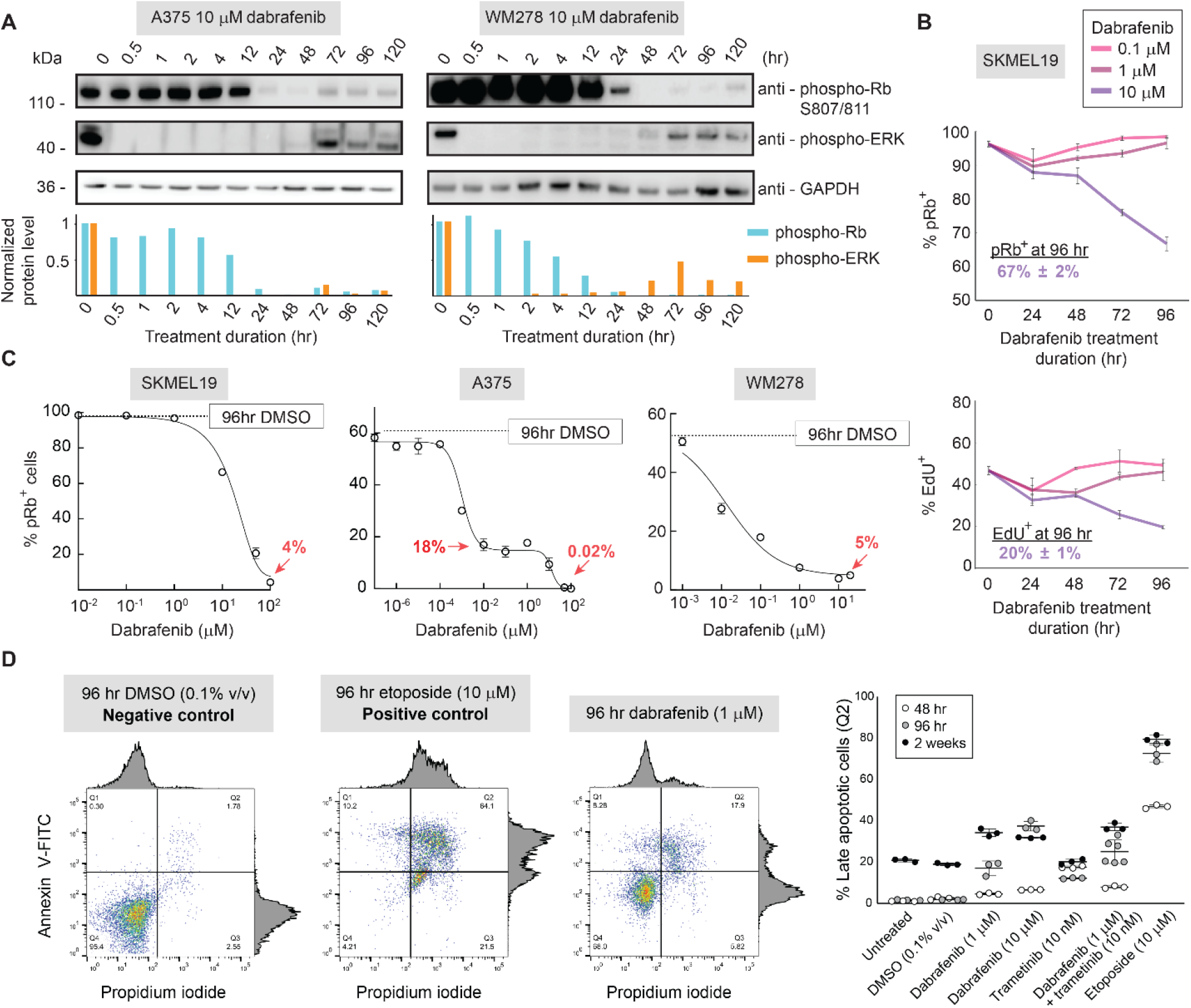
Dabrafenib treatment induces quiescence incompletely. (A) A375 (left) and WM278 (right) cells were treated with 10 μM dabrafenib for the indicated durations and the levels of phospho-ERK and phospho-Rb S807/811 were measured by western blot (top). Signals were quantified by normalizing first to GAPDH levels then to the untreated condition (bottom). (B) Quantification of percentage of pRb^+^ cells in SKMEL19 cells treated for indicated lengths of time with 0.1 μM, 1 μM, and 10 μM dabrafenib. The percentage of pRb^+^ cells with 10 μM dabrafenib at 96 hr is noted. Error bars: as mean ± std of 3 replicate wells. (C) SKMEL19, A375 and WM278 dose-response curves showing the percent pRb^+^ cells after 96 hr of dabrafenib treatment, determined by immunofluorescence quantification. For A375, the untreated 96 hr DMSO line falls at 60% (compared with 95% reported elsewhere in this study when cells were plated 24 hr before fixation) because 96 hr of unfettered growth on the plate results in partial contact inhibition. Error bars: as mean ± std of 3 replicate wells. (D) Apoptotic cell quantification by flow cytometric analyses of Annexin V-FITC and propidium iodide staining. DMSO bivariate plot is shown as a negative control for apoptosis; etoposide is shown as a positive control. Representative bivariate plot is shown for 96 hr of 1 μM dabrafenib. Quantified replicates of late apoptotic cells (Q2) in each treatment condition are shown for the indicated time points, with mean represented as a horizontal line. Error bars: as mean ± std of at least 3 replicate samples.

**Fig. S2.**
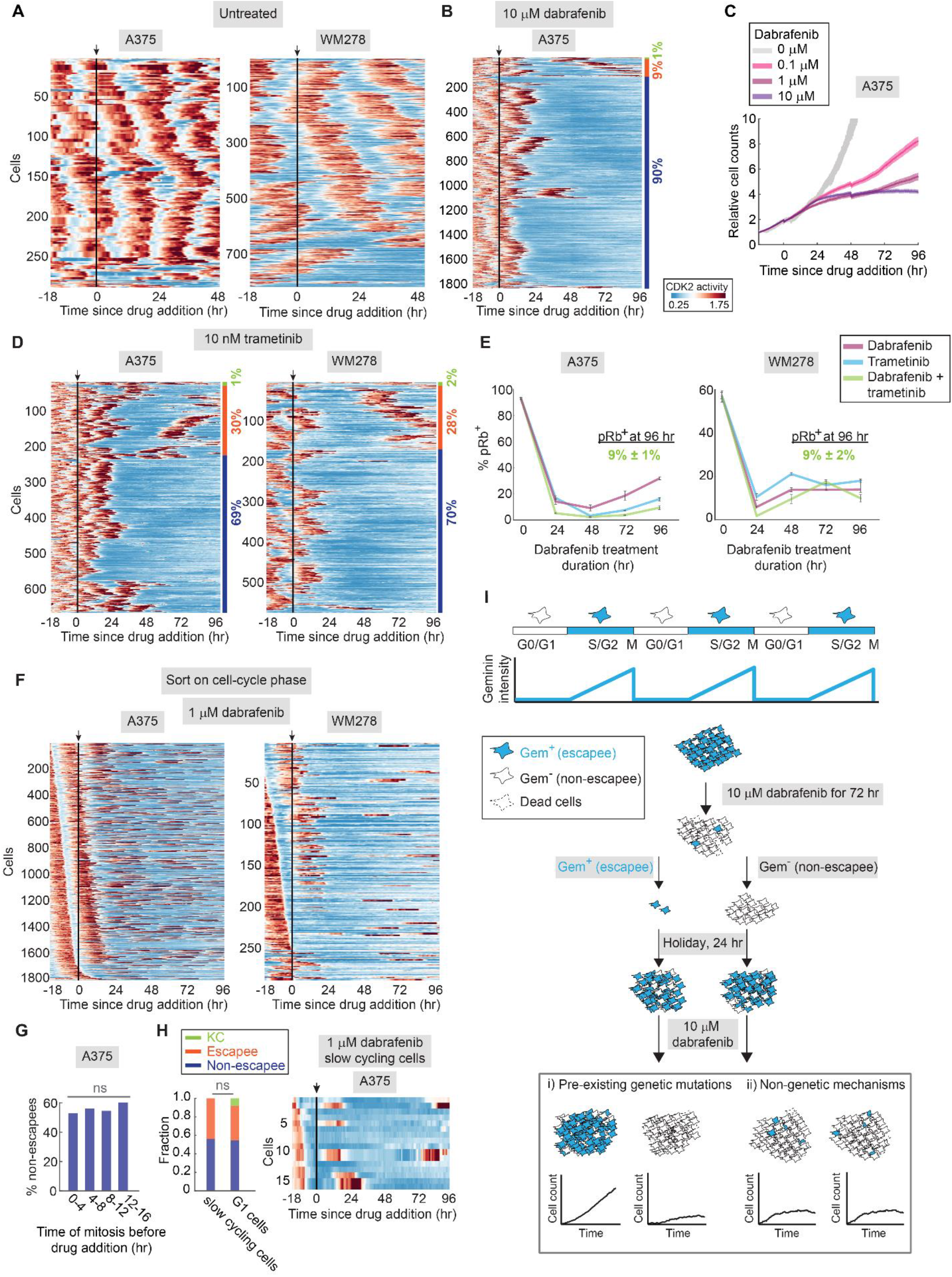
Cellular heterogeneity in drug response is revealed by live-cell imaging, and escapees do not arise from cells with pre-existing mutations. (A-B) Heatmap of single-cell CDK2 activity traces in untreated or 10 μM dabrafenib-treated A375 and WM278 cells. Each row represents the CDK2 activity in a single cell over time according to the colormap in which blue indicates low activity and red indicates high activity. Cells were first sorted by cell-cycle behavior and then sorted by similarity of CDK2 traces. The black line marks the time of media change or drug addition. Drug was refreshed at 48 hr. Apoptotic cells are not included in the heatmap. The percentages mark the proportion of cells with each behavior. KC, keep cycling. (C) Relative cell count over time in A375 cells treated with dabrafenib for 96 hr, as monitored by time-lapse microscopy. Error bar, 95% confidence interval. (D) Heatmap of single-cell CDK2 activity traces in 10 nM trametinib-treated A375 and WM278 cells. In addition to serving as a control for Fig. 1H, this heatmap also shows the existence of escapees upon treatment with a clinical MEK1/2 inhibitor. (E) Quantification of percentage of pRb^+^ cells in A375 and WM278 cells treated for the indicated durations with dabrafenib or trametinib alone or in combination. The percentage of pRb^+^ cells under the combined treatment at 96 hr is noted. Error bars: mean ± std of 3 replicate wells. (F) Replotting of data in Fig. 1G by sorting traces according to how long ago cells underwent mitosis prior to drug addition. (G) Bar plot of fraction of non-escapees in different cell-cycle phases at the time of 1 M dabrafenib addition. Cells were classified into different cell-cycle phases based on the time of mitosis before drug treatment. Then the fraction of non-escapees was calculated based on the cell behavior over the 96 hr drug treatment duration. For each category, a binomial test was performed to evaluate the difference between the reported fraction and the fraction in all cells (57%, Fig. 1G). (H) Percentage of keep cycling, escapee, and non-escapee cells between naturally slow-cycling cells and G1 cells. Cells were designated slow-cycling if they were in a CDK2^low^ quiescence for more than 6 hr prior to drug addition; 16 cells met this criterion. Cells were designated as G1 cells if CDK2 activity rose no more than 6 hr prior to drug addition; 625 cells met this criterion. Left panel: percentage of each behavior in slow-cycling and G1 cells treated with 1 μM dabrafenib. A binomial test was performed to compare the fraction of non-escapees in the two categories. Right panel: heatmap of CDK2 activity for 16 slow-cycling cells sorted according to how long ago cells underwent mitosis prior to drug addition, showing that these cells can still readily escape drug. (I) Schematic diagram of the drug holiday experimental setup described in the text.

**Fig. S3.**
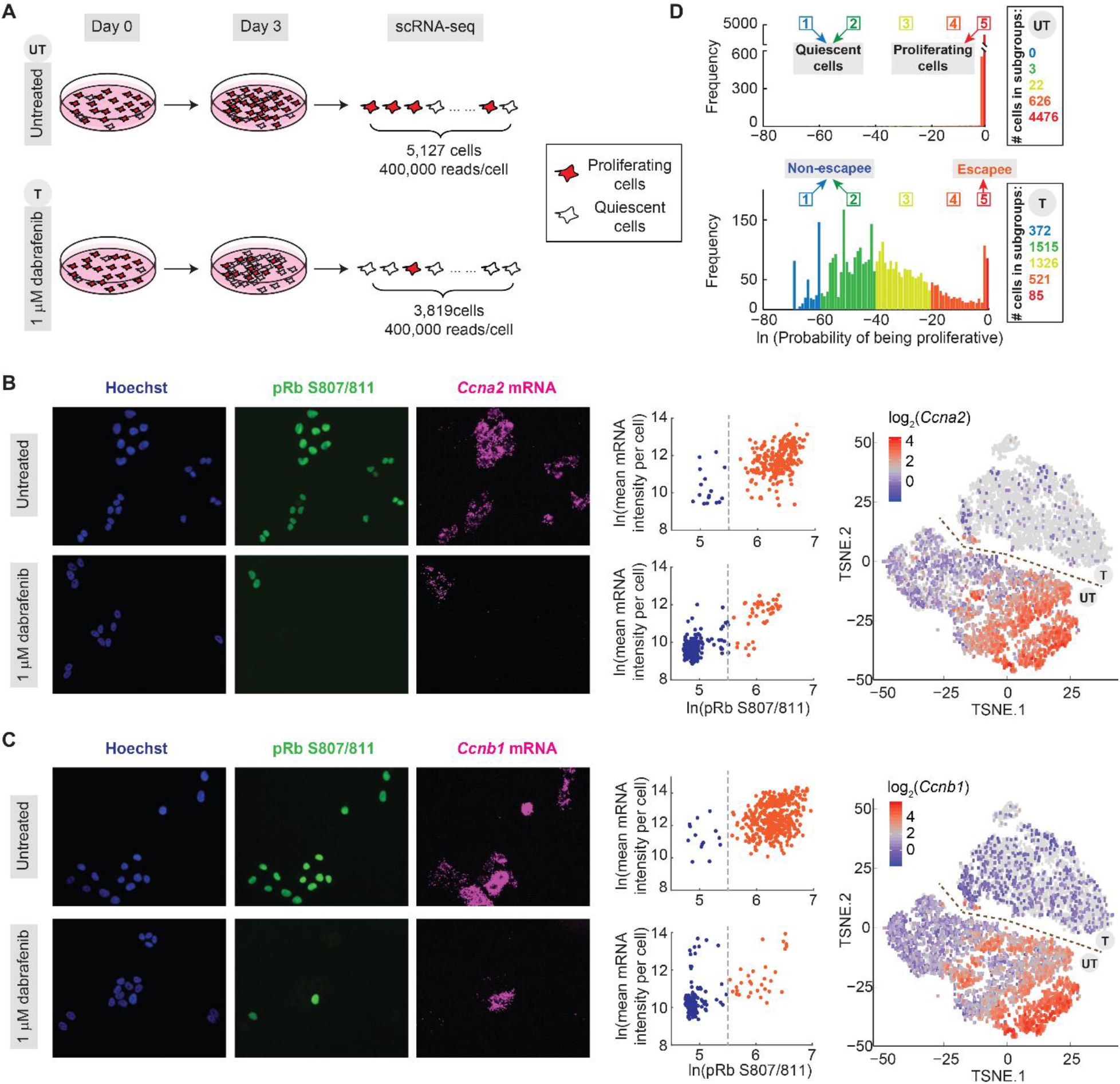
Calculation of proliferation probability based on 51 cell-cycle genes and validation of two of the genes. (A) Schematic of scRNA-seq experiment. (B-C) mRNA expression levels of *Ccna2* and *Ccnb1* in untreated A375 cells or in cells treated with 1 μM dabrafenib for 72 hr. Left: co-staining of phospho-Rb and the indicated mRNA; Middle: quantification of mRNA levels in pRb^+^ and pRb^−^ cells (each population pooled from 2 replicate wells); Right: tSNE plots based on scRNA-seq data showing increased expression of these two genes in the escapee subpopulation, visible as a small peninsula of red-shaded cells in the treated population. (D) Histograms of single-cell proliferation probability in untreated A375 cells (UT, top) and cells treated with 1 μM dabrafenib for 72 hr (T, bottom) conditions. Cells were placed into five categories based on their probability of proliferation (see Methods and Extended Data Table1). The number of cells in each category is shown on the right with colors matching their categories.

**Fig. S4.**
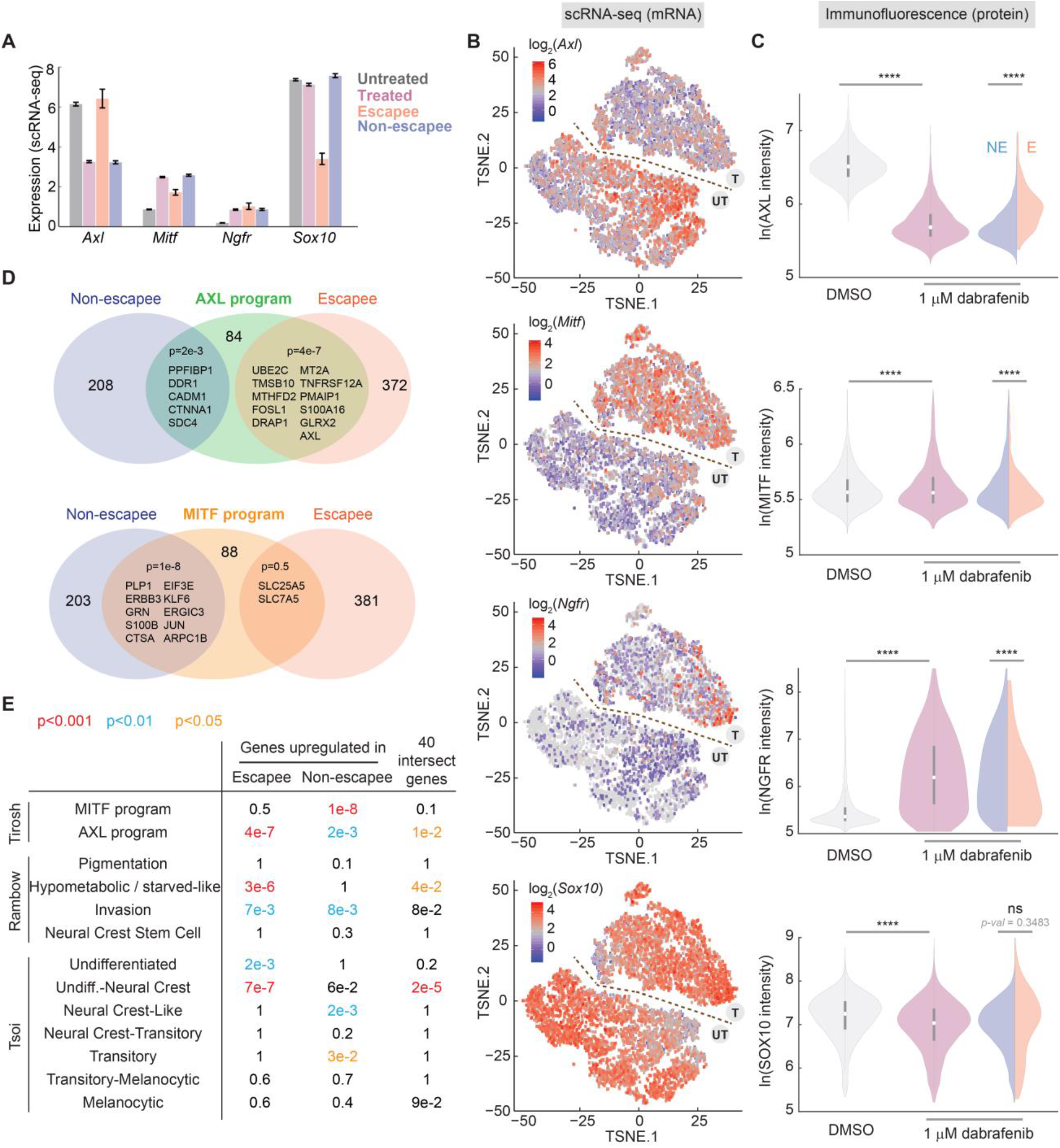
Escapees show an AXL^high^/MITF^low^ gene signature. (A) mRNA expression from scRNA-seq for *Axl, Mitf, Ngfr,* and *Sox10* in A375 cells. Untreated cells: all untreated cells from scRNA-seq; Treated cells: all treated cells from scRNA-seq; escapees: subgroup 5 in treated condition; non-escapees: subgroup 1 and 2 in treated condition. Bar plot indicates the mean ± SEM, where the SEM reflects technical noise and cell-to-cell variability. (B) Visualization of single-cell *Axl, Mitf, Ngfr,* and *Sox10* mRNA expression levels on the combined untreated and treated t-SNE plot. (C) Violin plot showing the AXL, MITF, NGFR, and SOX10 immunofluorescence signal in A375 cells treated with DMSO or 1 μM dabrafenib for 72 hr. Split violin plot indicates the protein level in dabrafenib-treated escapees (E) and non-escapees (NE), determined by co-staining with phospho-Rb (S807/811) in the case of MITF or phospho-Rb (S780) for the other markers. Each population value is pooled from 3 replicate wells. (D) Venn diagram showing the overlap of upregulated genes in escapees (subgroup 5 in treated condition) or non-escapees (subgroup 1 and 2 in treated condition) with the AXL or MITF program published in Tirosh *et al.* (*21*). Genes upregulated in escapees, n = 383; genes upregulated in non-escapees, n = 213; AXL-program: n = 100; MITF-program: n = 100. p-values were computed with the R package ‘GeneOverlap’ (Methods). (E) Comparison of upregulated genes in escapees or non-escapees with existing gene signatures. Comparison with Tirosh *et al.* was reproduced from (D). Sizes of gene signatures in Rambow *et al.* (*22*): pigmentation, n = 15; hypometabolic/starved-like, n = 27; invasion, n = 49; neural crest stem cell, n = 37. Sizes of gene signatures in Tsoi *et al.* (*23*): undifferentiation, n = 118; undiff.-neural crest, n = 106; neural crest-like, n = 66; neural crest-transitory, n = 25; transitory, n = 29; transitory-melanocytic, n = 125; melanocytic, n = 62.

**Fig. S5.**
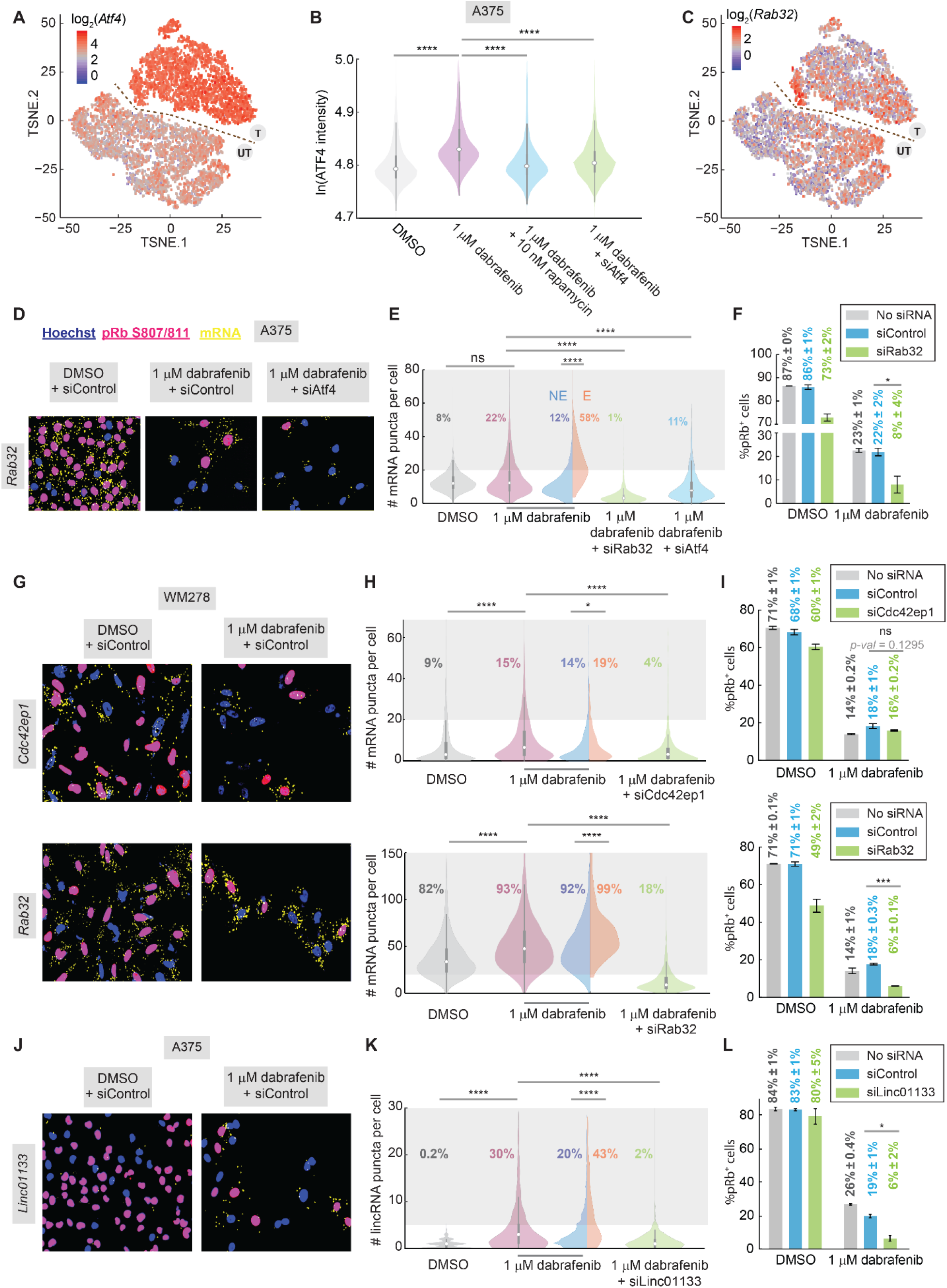
ATF4 target genes promote escape from dabrafenib in A375 and WM278 cells. (A) Visualization of single-cell ATF4 mRNA levels on the combined t-SNE plot, showing homogenous upregulation of ATF4 mRNA in all treated cells. (B) Violin plots showing ATF4 protein levels by immunofluorescence in A375 cells treated as indicated. ATF4 expression is significantly induced by dabrafenib but repressed by addition of the mTORC1 inhibitor rapamycin or siATF4. Each population value is pooled from 2 replicate wells. (C) Visualization of single-cell *Rab32* mRNA levels on the combined t-SNE plot showing enrichment in the escapee subpopulation, visible as a small peninsula of red-shaded cells in the treated population. (D) Representative RNA-FISH images for *Rab32* together with phospho-Rb (S807/811) and Hoechst staining, for A375 cells treated for 72 hr with DMSO and siControl (left panel), 1 μM dabrafenib and siControl (middle panel), or 1 μM dabrafenib and siRNA against ATF4 (right panel). (E) Quantification of number of mRNA puncta for *Rab32* in each condition indicated. The percentage of cells that have > 20 mRNA puncta is indicated on the plot. Each population value is pooled from 2 replicate wells. (F) The percentage of pRb^+^ A375 cells under treatment with DMSO or 1 μM dabrafenib and either no siRNA (grey), control siRNA (blue) or siRNA against *Rab32* (green). Error bars: mean ± std of 4 replicate wells. (G) Representative images of WM278 cells stained for *Cdc42ep1* or *Rab32* mRNA, phospho-Rb, and Hoechst, for the indicated 72 hr treatment conditions. (H) Quantification of number of mRNA puncta in WM278 cells for *Cdc42ep1* (upper panel) and *Rab32* (lower panel) in each condition indicated. The percentage of cells that have > 20 mRNA puncta for *Cdc42ep1* or *Rab32* are indicated on the plot of each condition. Each population value is pooled from 2 replicate wells. (I) The percentage of pRb^+^ WM278 cells after 72 hr treatment with DMSO or 1 μM dabrafenib and either no siRNA (grey), control siRNA (blue) or siRNA against *Cdc42ep1*(upper panel) or *Rab32* (lower panel) (green). Error bars: mean ± std of 4 replicate wells. (J-L), Same analysis as (G-I), but for *Linc01133* in A375 cells. The percentage of cells that have > 5 mRNA puncta for *Linc01133* is indicated on the plot of each condition. For the DMSO-treated violin plot in (K), most cells have zero puncta and only a few cells have low integer numbers of puncta, thus the violin plot appears staggered at the integer values.

**Fig. S6.**
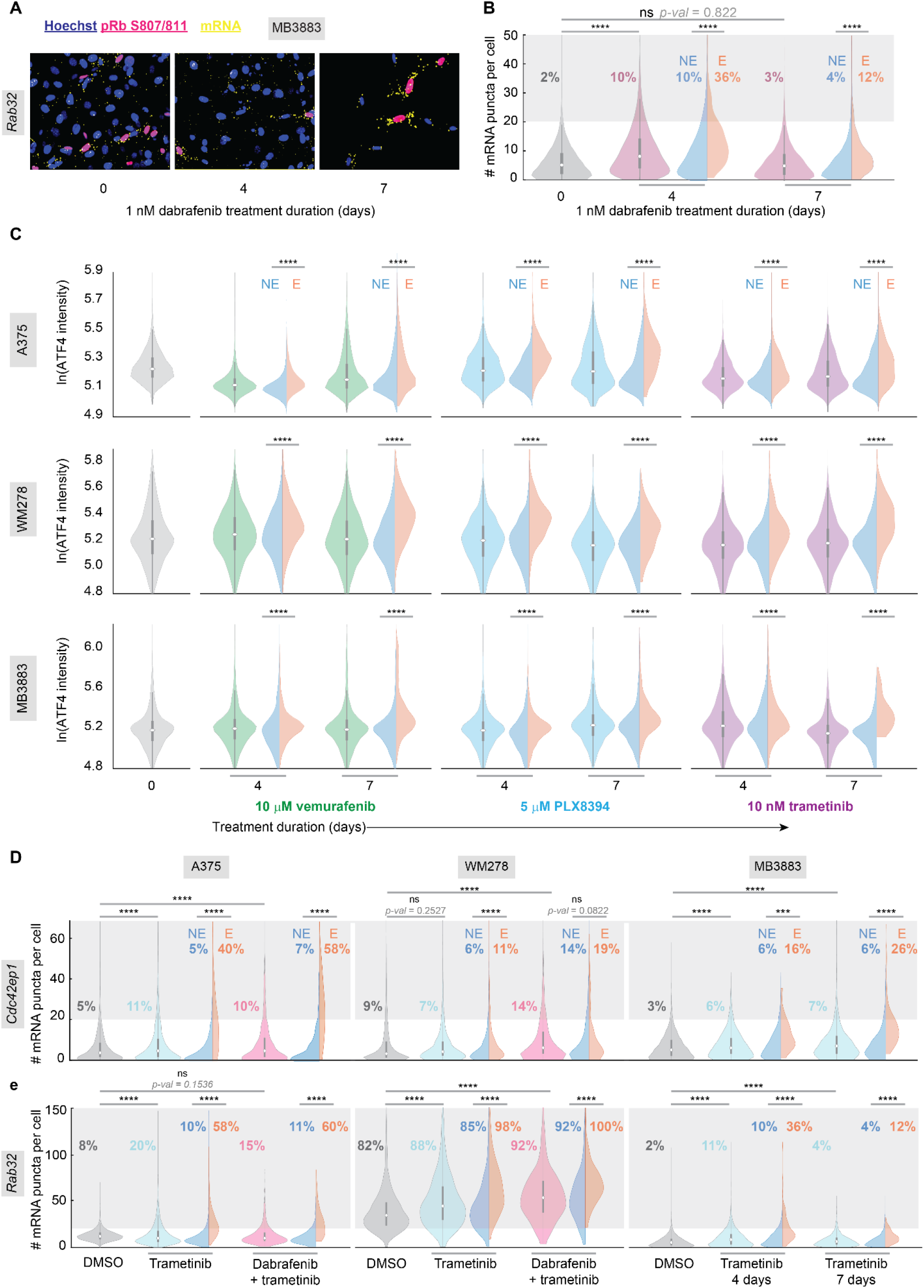
The escapee phenotype is observed with multiple MAPK pathway inhibitors and also occurs in *ex vivo* patient biopsies. (A) Representative RNA-FISH images for *Rab32* together with pRb (S807/811) and Hoechst staining, for MB3883 patient cells cultured in 1 nM dabrafenib for 0, 4, and 7 days. (B) Violin plot of the conditions in (A) showing the number of *Rab32* mRNA puncta in MB3883 cells. Split violin plot indicates the number of puncta in dabrafenib-treated escapees (E) and non-escapees (NE). The percentage of cells that have > 20 mRNA puncta is indicated on the plot of each condition. Each population value is pooled from 2 replicate wells. (C) Violin plots of nuclear ATF4 protein intensity in A375, WM278, and MB3883 cells treated with the indicated drugs for 0, 4, or 7 days. Each population value is pooled from 4 replicate wells. (D-E), Violin plots showing the number of *Cdc42ep1* (D) or *Rab32* (E) mRNA puncta in A375 and WM278 cells treated with DMSO, 10 nM trametinib, or 1 μM dabrafenib plus 10 nM trametinib for 72hr. Far-right section shows the number of *Cdc42ep1* (D) or *Rab32* (E) mRNA puncta in MB3883 cells treated for 0, 4 or 7 days with 1 nM trametinib. The percentage of cells that have > 20 mRNA puncta for each gene is indicated on the plot. Each population value is pooled from 2 replicate wells.

**Fig. S7.**
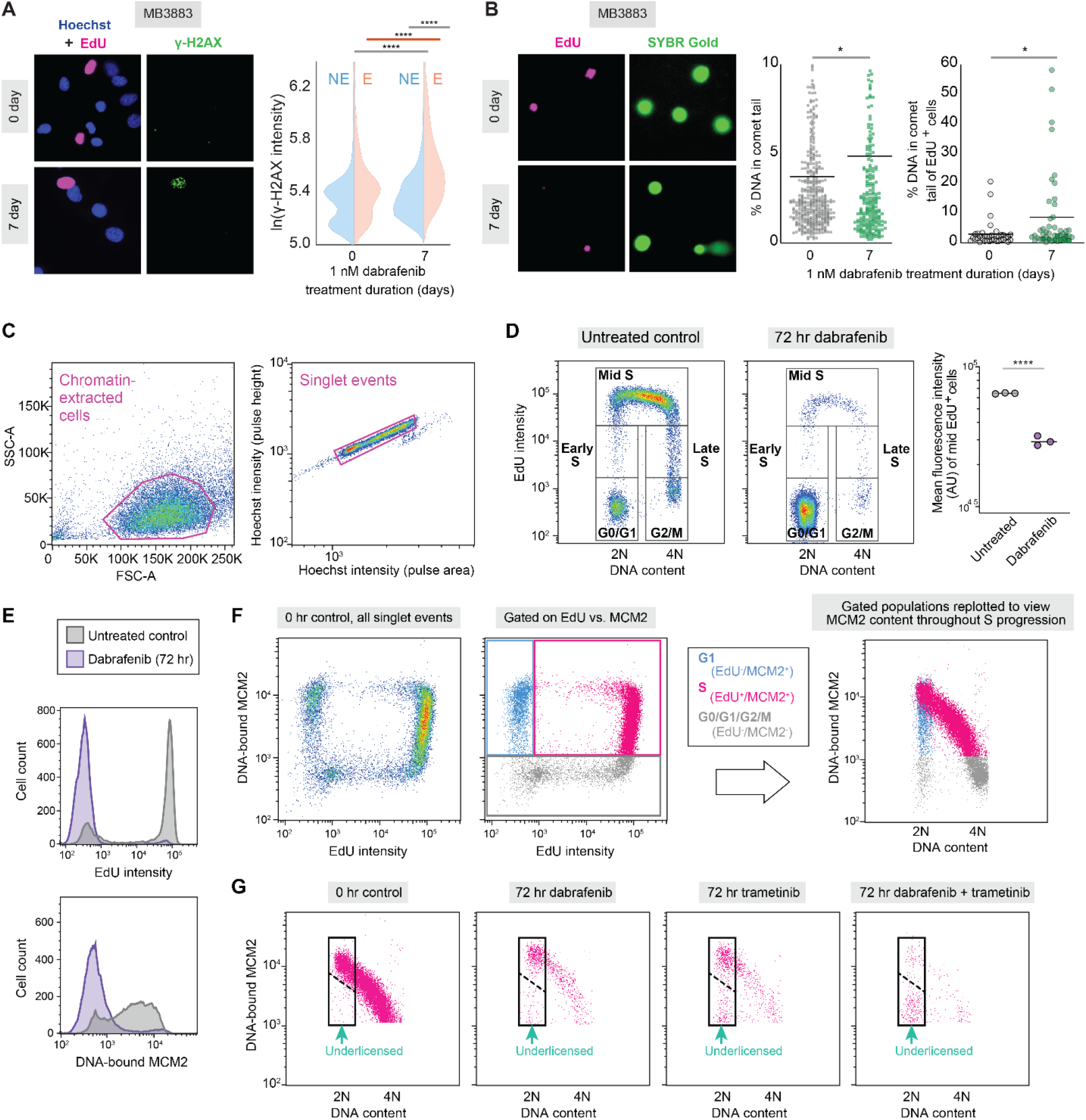
Significant DNA damage is also observed in escapees detected in *ex vivo* patient cultures, and cells cycling in the presence of dabrafenib and trametinib do not properly load MCM2 on DNA when attempting to license origins of replication. (A) Representative images of 1 nM dabrafenib-treated MB3883 patient cells stained for EdU incorporation and γ-H2AX. Quantifications of the γ-H2AX puncta for escapees (E) and non-escapees (NE) are plotted as split violins. Each population value is pooled from 6 replicate wells. (B) Neutral comet assay analysis in 1 nM dabrafenib-treated MB3883 patient cells. Images show gels co-stained for SYBR Gold to mark DNA and EdU to mark cells in S phase. Plots show all cells (left) or only EdU^+^ cells (right) in which the percent of DNA in each comet tail was measured, with mean values indicated on the plots as horizontal lines. Each population value is pooled from 2 biological comet slide replicates. (C) Gating scheme to identify single chromatin-extracted cells by flow cytometry. (D) Scatter plot of DNA synthesis rate (EdU incorporation, 30 min pulse) vs. DNA content (integrated Hoechst signal) in untreated A375 cells to identify cell-cycle phases. The same gates are propagated to the plot of dabrafenib-treated cells. Right-most plot shows mean fluorescence intensity of all cells in mid S phase (3 replicate samples). (E) Representative histograms of DNA synthesis rate (EdU intensity) and chromatin-associated MCM2 for untreated and dabrafenib-treated cells. (F) Gating scheme (as in ref. *31*) used to identify cell-cycle phases by EdU and chromatin-bound MCM2 signal intensity in untreated cells. Cells that are both EdU^+^ and chromatin-bound MCM2^+^ are shaded pink. (G) Scatter plot of EdU^+^ and chromatin-bound MCM2^+^ cells gated as in (F) for the indicated treatment conditions (pooled from 3 replicate samples). Untreated cells showing normal licensing of replication origins are used to draw the initial gate (left-most plot), and this gate is then propagated to the treated conditions. Cells falling below the dashed line in this gate are under-licensed. The percentage of under-licensed cells (below dashed line) out of all early S cells (entire rectangle) is reported in Fig. 4 for 3 replicates.

**Fig. S8.**
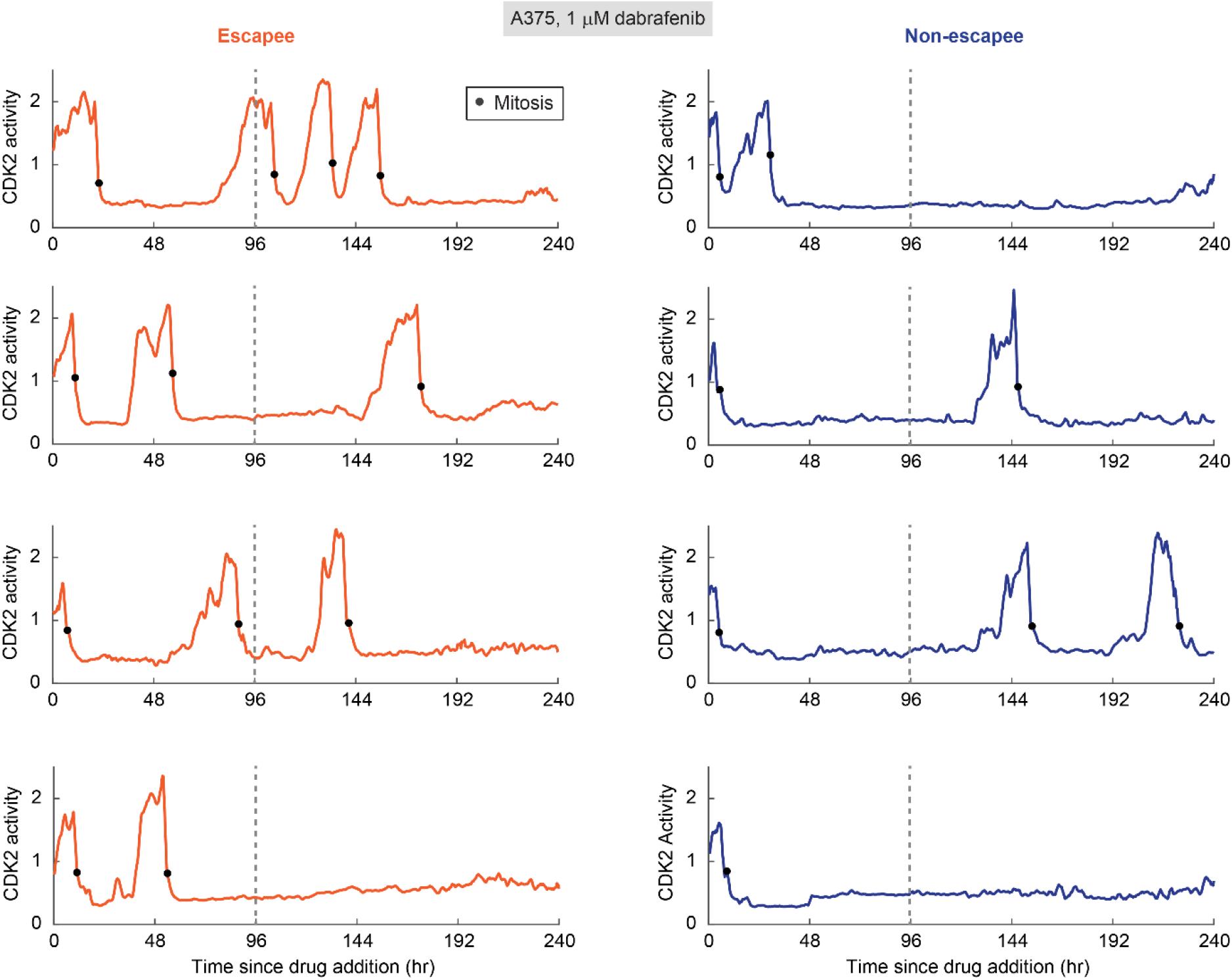
Escapees outgrow non-escapees over extended treatment. Sample single-cell CDK2 activity traces from a 10-day movie of A375 cells treated with 1 μM dabrafenib at the start of the movie. Escapees and non-escapees are defined by their behavior during the first 96 hr of filming (dashed line), as in Fig.4E. Black dots mark mitoses.

